# Identification of polyketide biosynthetic gene clusters that harbor self-resistance target genes

**DOI:** 10.1101/2020.06.01.128595

**Authors:** Gergana A Vandova, Aleksandra Nivina, Chaitan Khosla, Ronald W Davis, Curt R Fisher, Maureen E Hillenmeyer

## Abstract

**Background:** Polyketide secondary metabolites have been a rich source of antibiotic discovery for decades. Thousands of novel polyketide synthase (PKS) gene clusters have been identified in recent years with advances in DNA sequencing. However, experimental characterization of novel and useful PKS activities remains complicated. As a result, computational tools to analyze sequence data are essential to identify and prioritize potentially novel PKS activities. Here we exploit the concept of genetically-encoded self-resistance to identify and rank biosynthetic gene clusters for their potential to encode novel antibiotics.

**Results:** To identify PKS genes that are likely to produce an antibacterial compound, we developed an automated method to identify and catalog clusters that harbor potential self-resistance genes. We manually curated a list of known self-resistance genes and searched all NCBI genome databases for homologs of these self-resistance genes in biosynthetic gene clusters. The algorithm takes into account (1) the distance of the potential self-resistance gene to a core enzyme in the biosynthetic gene cluster; (2) the presence of a duplicated housekeeping copy of the self-resistance gene; (3) the presence of close homologs of the biosynthetic gene cluster in diverse species also harboring the putative self-resistance gene; (4) evidence for coevolution of the self-resistance gene and core biosynthetic gene; and (5) self-resistance gene ubiquity. We generated a catalog of 190 unique PKS clusters whose products likely target known enzymes of antibacterial importance. We also present an expanded set of putative self-resistance genes that may be useful in identifying small molecules active against novel microbial targets.

**Conclusions:** We developed a bioinformatic approach to identify and rank biosynthetic gene clusters that likely harbor self-resistance genes and may produce compounds with antibacterial properties. We compiled a list of putative self-resistance genes for novel antibacterial targets, and of orphan PKS clusters harboring these targets. These catalogues are a resource for discovery of novel antibiotics.

## Introduction

DNA sequencing has revealed thousands of new biosynthetic gene clusters (BGCs) in microbial genomes. Many BGCs encode assembly-line polyketide synthases (PKSs), large multifunctional enzymes that produce biologically active natural products, many of them used in the clinic. As of 2013, NCBI nucleotide databases contained ∼900 assembly-line PKSs - a number that has grown to ∼3,500 by 2018^1,2^. Most of them are “orphan” or “cryptic”, meaning these gene clusters have not been characterized, and any molecules they might produce are uncharacterized. Moreover, 544 type II PKSs were cataloged as of 2015, most of which are also orphan^3^. However, experimental characterization of these gene clusters is complicated, labor-intensive, and with low success rate^4^. Therefore, computational tools that identify BGCs likely to encode novel activities are needed. There are several automated genome mining tools that predict the presence of a BGC in a bacterial, fungal, or plant genome: antiSMASH^5^, plantiSMASH^6^, ClusterFinder^7^, SMURF^8^, ClustScan^9^, CLUSEAN^10^. However, prioritizing novel and useful PKS activities remains a bottleneck. A key question is how to use available sequence data to prioritize BGCs likely to encode compounds with novel bioactivities.

In order to prioritize gene clusters, we exploit the concept of self-resistance^11-13^. To avoid self-toxicity, these antibiotic-producing microorganisms sometimes encode a resistant copy of the target gene. Often, these self-resistance genes are co-localized and co-expressed with the antibiotic biosynthetic genes. For example, the biosynthetic gene cluster of the fatty acid synthase inhibitor thiotetronic acid contains a resistant copy of the fatty acid synthase gene (*fabB*/*F*)^12^. This work focuses solely on self-resistance genes that are resistant copies of the targets of the produced compound (hereafter self-resistance genes). We hypothesized that by searching for bacterial BGCs that harbor a bacterial self-resistance gene, we could increase the chance of identifying clusters that produce molecules with antibacterial activity. We thus developed an automated method to identify and catalog clusters that harbor a potential self-resistance gene. A similar idea was used by the Antibiotic Resistant Target Seeker (ARTS) web tool, which allows for rapid identification of known self-resistance mechanisms based on BGC proximity, presence of an extra copy of the gene in actinobacterial genomes, and phylogeny-based analyses in Actinobacteria^14^. Our approach is complementary to the ARTS tool and generalizes to BGCs from any bacterial species. Similar to ARTS, we used evidence of gene duplication and proximity to the BGC as screening criteria. We differ in use of a more stringent distance cutoff between a core KS domain and the putative self-resistance gene, and we developed an additional criterion that reflects the coevolution of the proximal KS domain and the putative self-resistance gene. We validated the algorithm’s performance on a manually-curated list of gene clusters known to harbor a self-resistance gene. We applied our approach to detect new gene clusters harboring these genes as putative self-resistance genes. We propose that these clusters should be prioritized for further characterization.

We sought to generalize our approach to identify other gene families that might provide self-resistance in BGCs. We present an expanded set of putative self-resistance genes that may be useful in identifying BGCs that encode small molecules with novel microbial targets. We applied our method to search for clusters harboring these genes, and present the highly ranked orphan PKS clusters most likely to produce antibacterial compounds. Our approach predicts the potential targets of those compounds, and as such could be a useful tool in biosynthetic cluster prospecting.

## Materials and Methods

### Curation of a reference set of experimentally validated self-resistance genes

In order to identify bacterial type I PKS BGCs harboring a self-resistance copy of a target gene, we curated a list of known antibacterial self-resistance genes co-localized with the type I PKS BGCs in the Minimum Information About a Biosynthetic Gene Cluster (MIBiG) database^15^. All 1816 cluster files from MIBiG 1.4 database were parsed to find putative targets for each MIBiG entry (“Molecular Target” field in the MIBiG database). UniProt database was searched for a set of proteins matching this description. We made a BLAST database for each cluster with all of the proteins in that cluster. For each protein in the set of UniProt protein sequences, we used BLASTP to identify a homolog in the cluster. We filtered the BLASTP hits using identity threshold of 50% and “Molecular Target” MIBiG field as “Antibacterial” or “Cytotoxic”. For any given hit from the BLAST search, we searched the literature for experimental evidence of self-resistance of that gene in that particular cluster. Based on these results and literature search, we identified 18 clusters harboring one of 14 experimentally validated self-resistance protein families. These clusters biosynthesize polyketides, non-ribosomal peptides and their hybrids, terpenes, ribosomally synthesized and post-translationally modified peptides (RiPPs) and aminocumarins. Table 1 lists the number of clusters harboring each of these 14 self-resistance gene families.

**Table 1.** Experimentally validated gene clusters harboring a self-resistance gene. These natural products are encoded by gene clusters (**MIBiG ID)** that harbor an experimentally-validated self-resistance gene. Compounds denoted with (*) are encoded by a closely-related gene cluster harboring the same putative self-resistance gene, but have not yet been experimentally validated to provide the self-resistance mechanism.

| N | Self-resistance Protein | Natural Product |  | MIBiG ID | Class | Target Pathway | Distance to Core Enzyme | Refs. |
| --- | --- | --- | --- | --- | --- | --- | --- | --- |
| 1 | 3-oxoacyl-[acyl-carrier-protein] synthase 1 (FabB/F) | 1 | thiolactomycin | BGC0001237 | PKS | fatty acid biosynthesis | 1.3 kB | <sup>12</sup> |
|  |  | 2 | thiotetramide | BGC0001236 | PKS | fatty acid biosynthesis | 1.4 kB | <sup>12</sup> |
|  |  | 3 | platensimycin / platencin | BGC0001140/<br>BGC0001156 | terpene | fatty acid biosynthesis | 13 kB | <sup>44</sup> |
| 2 | Acetyl CoA carboxylase | 4 | andrimid | BGC0000956 | PKS/NRPS | fatty acid biosynthesis | 5.1 kB | <sup>45</sup> |
| 3 | Beta proteasome subunit | 5 | salinosporamide | BGC0001041 | PKS/NRPS | proteasome inhibitor | 7.7 kB | <sup>40</sup> |
|  |  | 6 | cinnabaramide* | BGC0000971 | PKS/NRPS | proteasome inhibitor | 0.6 kB | <sup>46</sup> |
|  |  | 7 | clarepoxcin* | BGC0001203 | PKS/NRPS | proteasome inhibitor | 8.9 kB | <sup>47</sup> |
|  |  | 8 | eponemycin * | BGC0000345 | PKS/NRPS | proteasome inhibitor | 4.1 kB | <sup>48</sup> |
| 4 | DNA polymerase sliding clamp | 9 | griselimycin | BGC0001414 | NRPS | DNA replication | 0.7 kB | <sup>49</sup> |
| 5 | Elongation factor Tu | 10 | GE2270 | BGC0001155 | RiPP | protein synthesis | N/A | <sup>50</sup> |
|  |  | 11 | GE37468* | BGC0000605 | RiPP | protein synthesis | N/A | <sup>51</sup> |
|  |  | 12 | TP-1161* | BGC0000615 | RiPP | protein synthesis | N/A | <sup>52</sup> |
|  |  | 13 | kirrothricin* | N/A | PKS/NRPS | protein synthesis | N/A | <sup>53</sup> |
|  |  | 14 | efrotomycin* | N/A | PKS/NRPS | protein synthesis | N/A | <sup>53</sup> |
| 6 | Enoyl-[acyl-carrier-protein] reductase [NADH] FabI | 15 | kalimantacin / batumin | BGC0001099 | PKS/NRPS | fatty acid biosynthesis | 4.8 kB | <sup>54</sup> |
| 7 | Gyrase B | 16 | microcin B17 | BGC0000568 | RiPP | DNA replication | N/A | <sup>55</sup> |
|  |  | 17 | novobiocin | BGC0000834 | aminocoumarin | DNA replication | N/A | <sup>56</sup> |
|  |  | 18 | clorobiocin | BGC0000832 | aminocoumarin | DNA replication | N/A | <sup>57</sup> |
|  |  | 19 | coumermycin A1 | BGC0000833 | aminocoumarin | DNA replication | N/A | <sup>57</sup> |
| 8 | Isoleucyl-tRNA synthetase | 20 | mupirocin | BGC0000182 | PKS | protein synthesis | 4.6 kB | <sup>58</sup> |
|  |  | 21 | thiomarinol | BGC0001115 | PKS/NRPS | protein synthesis | 4.5 kB | <sup>59</sup> |
| 9 | Leucil-tRNA synthase | 22 | agrocin 84 | N/A | Other | protein synthesis | N/A | <sup>60</sup> |
| 10 | Methionine aminopeptidase | 23 | bengamide | N/A | PKS/NRPS | protein synthesis | 4.1 kB | <sup>61</sup> |
| 11 | Ornithine carbamoyl transferase | 24 | phaseolotoxin | BGC0000919 | Other | amino acid synthesis | N/A | <sup>62</sup> |
| 12 | Seryl-tRNA synthetase | 25 | albomycin | N/A | Sideromycin | protein synthesis | 4.4 kB | <sup>63</sup> |
| 13 | Threonyl-tRNA synthetase | 26 | borrelidin | BGC0000031 | PKS | protein synthesis | 14.5 kB | <sup>64</sup> |
| 14 | Tryptophanyl-tRNA synthase | 27 | indolmycin | BGC0001206 | Other | Protein synthesis | N/A | <sup>65</sup> |

### Identifying Type I PKS clusters harboring a putative self-resistance gene

#### Step 1. Performing BLAST search for KS homologs

All NCBI nucleotide and genome databases were searched for KS homologs using TBALSTN (Figure 1). 8 diverse KS sequences from modular Type I PKS (erythromycin), cis-AT PKS/NRPS (curacin, epothilone, guadinomine, rapamycin) and trans-AT PKS/NRPS (leinamycin, disorazol, chivosazol) clusters were used as query sequences against the major NCBI nucleotide and genome databases (nt, refseq_genomic, other_genomic, env_nt, patnt, htgs, tsa_nt, sts, gss, est_others updated in April 2018 and wgs updated in January 2016), as well as the Bacterial Assemblies database updated in October 2018.

**Figure 1.**
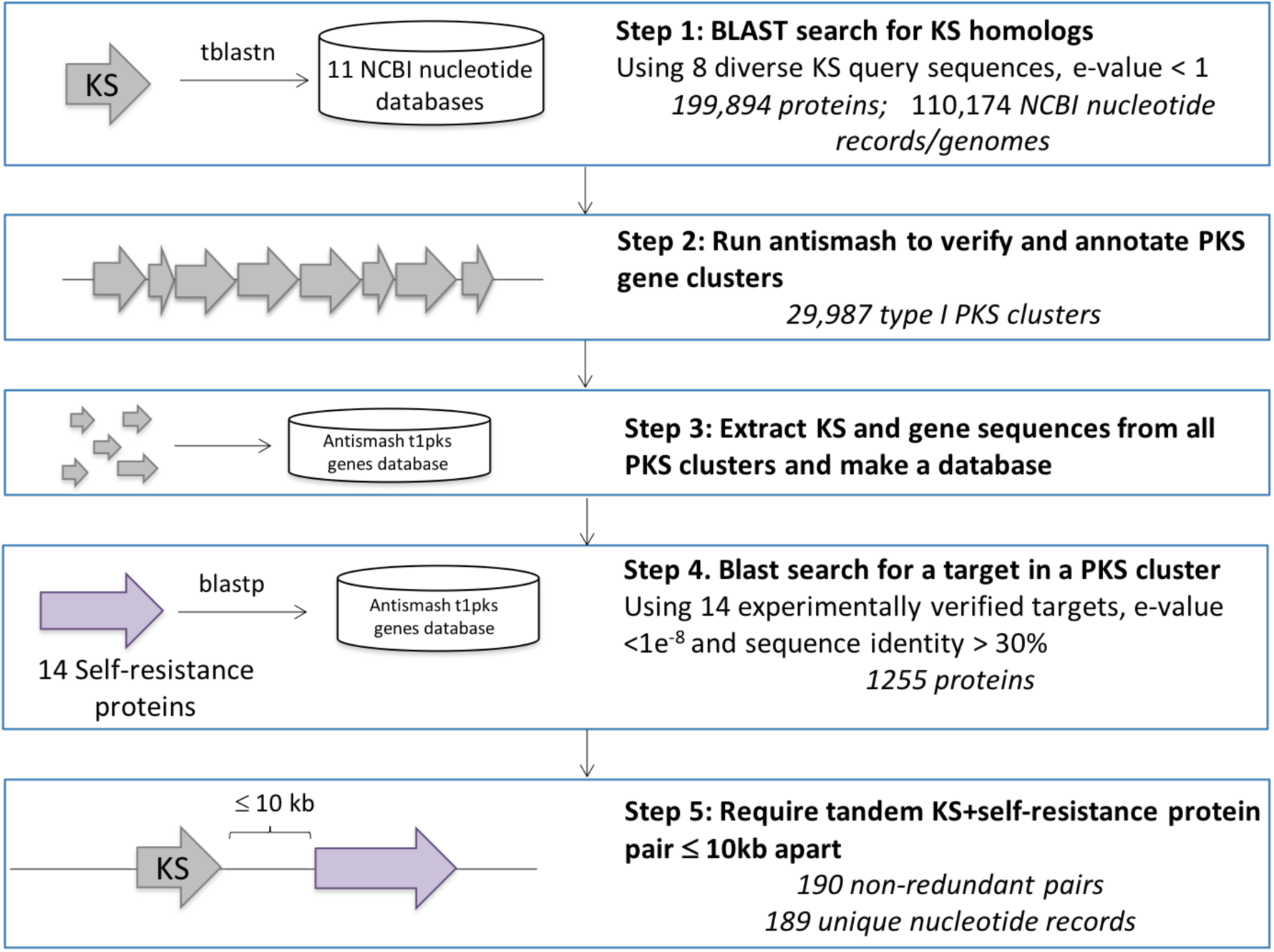
Identifying PKS clusters harboring a putative self-resistant target gene.

An initial relaxed blast search (e-value < 1) identified 199,894 non-redundant protein sequences (99% identity cutoff) contained in 110,174 unique NCBI nucleotide records/genomes.

#### Step 2. Running antiSMASH

antiSMASH 4.0^16^ with minimal functionalities was run on all 79,053 non-redundant NCBI records over 10,00bp in length, including 26,573 assembly records. To speed up computation time, assembly records were split into smaller ones by extracting sequence 150 kB upstream and downstream of a KS domain and running antiSMASH in parallel.

#### Step 3: Generating a database of proteins found in Type I PKS clusters

antiSMASH identified 29,987 clusters as Type I PKS, which constituted our cluster database. We extracted all proteins from these clusters, resulting in a database of 908,532 protein sequences. We also extracted all KS domains from these clusters and constructed a KS database.

#### Step 4. Performing BLAST search for putative self-resistance genes in PKS clusters

We searched the protein database for 14 experimentally validated self-resistance proteins (Table 1) using BLASTP with a relaxed e-value <1, which resulted in 4,404 similar proteins. Filtering blast hits by e-value <1e^-8^ and identity > 0.3 (except for FabB/F self-resistance proteins, for which we used identity cutoff of 0.6 to be able to distinguish between a KS and a FabB/F since they share high sequence identity) reduced the set to 1255 similar proteins. The number of copies of the gene per NCBI record was counted (to later determine the presence of a second, putatively-housekeeping copy of the self-resistance gene), and the status of the record (complete genome vs. incomplete) was noted.

#### Step 5. Identifying putative self-resistance genes in proximity to KS domains

Most experimentally characterized self-resistance genes (Table 1) are located within 10 kB of a core KS domain, with the exception of *fabB/F* in platencimycin BGC (13 kB away) and threonyl-tRNA synthetase in borrelidin BGC (14.5 kB). We set an initial threshold for the maximum allowed distance between a KS domain and a putative self-resistance gene at 10 kB. This filter reduced the set of 1255 putative self-resistance genes to 489 in 465 clusters. For additional analyses, we compared with threshold of ≤ 5 kB.

### Phylogenetic analyses

To remove redundant gene clusters, we selected a threshold of 90% KS protein sequence identity, which resulted in 190 gene clusters with putative self-resistance genes in 189 unique NCBI nucleotide records. Multiple sequence alignment of KS domains located in proximity to a putative self-resistance gene was performed using MAFFT^17^ and phylogenetic tree was generated using FastTree 2^18^. The tree was visualized using the APE package in R^19^. The tree was rooted on *E. coli* FabB/F.

### Housekeeping copy of the putative self-resistance gene

To identify whether a genomic contig contained an additional (putatively housekeeping) copy of a putative self-resistance gene, we used the 14 self-resistance protein sequences as queries in BLASTP search of all NCBI records harboring clusters with putative self-resistance genes. We used e-value cutoff of 1e^-8^ and identity threshold of 0.3 and FabB/F identity threshold of 0.6 (to be able to distinguish between a KS and a FabB/F since they share high sequence identity) to exclude false positive hits of KS domains. The number of copies per NCBI record was counted and the status of the record (complete genome vs. incomplete) was noted.

### Coevolution

For clusters harboring the same type of putative self-resistance gene, we defined a coevolution score between the protein sequences of self-resistance proteins and proximal KS domains. For each pair of such clusters, we visualized the amino acid sequence identity of the putative self-resistance gene vs. the amino acid sequence identity of the proximal KS domains, using either a ≤ 5 kB or a ≤ 10 kB distance threshold. We excluded KSs shorter than 200 nucleotides (67 amino acids) to avoid misannotated KSs, as most known KS genes are > 200 nucleotides in length. We fist plotted pairwise identities from the positive set of gene clusters with predicted self-resistance gene. (For clarexpoxin, eponemycin, cinnabaramide: a homolog of the known target gene is co-localized with a cluster, but it was not experimentally validated to be the self-resistance gene.) We calculated a Spearman correlation coefficient for each putative self-resistance protein.

### Ranking scheme

We devised a scoring scheme to rank the identified Type I PKS clusters harboring a putative self-resistance gene, based on several parameters: (1) distance of self-resistance gene to the closest KS domain; (2) presence of homologs in diverse species; (3) presence of a housekeeping copy of the self-resistance gene; (4) coevolution score based on the Spearman correlation coefficient; (5) self-resistance gene ubiquity, or the number of genomes in which a putative self-resistance gene has been identified: a high number of occurrences would increase the false discovery rate and vise-versa.

To select a threshold for defining homologous gene clusters, we manually examined pairwise amino acid sequence identities of KS domains from clusters known to be homologous. Niddamycin and spiramycin produce identical compounds and their Loading Module KS pairwise sequence identity is 82%. Tylactone and chalcomycin produce closely related compounds and their Loading Module KS is 71% identical. We therefore chose a threshold for homologous clusters of 80% amino acid identity.

Detailed description of the scores for each parameter is given in Table S1.

### Availability of data and materials

All data supporting the conclusions of this article is published online at http://gvandova.com/publish/Data/. Python code to generate the datasets and reproduce the analyses in this article are available online at https://github.com/GerganaVandova/TargetMining.

## Results and Discussion

### Curation of a reference set of 14 experimentally validated self-resistance genes in 18 BGCs

In order to identify bacterial type I PKS BGCs harboring a self-resistance copy of the target gene, we curated a list of known antibacterial self-resistance genes co-localized with the type I PKS BGCs by (a) automated mining of annotated gene clusters in the MIBiG database, and (b) manual literature search. We identified 18 clusters encoding one of 14 experimentally validated self-resistance proteins (Table 1). These clusters produce polyketides, non-ribosomal peptides and their hybrids, terpenes, ribosomally synthesized and post-translationally modified peptides (RiPPs) and aminocoumarins.

Of all BGCs deposited in MIBiG, 176 produce compounds annotated to have antibacterial activity. At least 15 of 176 BGCs harbor modified target genes that have been experimentally validated to be the self-resistance gene. (3 of the identified 18 clusters are currently not deposited in MIBiG). Additionally, 7 BGCs harbor putative self-resistance genes that have not been experimentally validated, but are likely providing the self-resistance mechanism of the producer organisms, because a homolog of the known target protein of the polyketide is encoded in the PKS gene cluster. 9 of these 18 gene clusters are type I PKSs or type I PKS hybrids. We hypothesize that more than 176 clusters produce antibiotics, but for most BGCs in MIBiG, the activity of the biosynthesized molecule is not annotated. Likely only a fraction of BGCs encoding compounds with antibacterial activity have been characterized and only a fraction of the self-resistance genes identified.

It is unknown how often microbes employ the self-resistance mechanism through target modification. Other resistance mechanisms such as drug efflux or compound modification are also employed^20^. Self-resistance genes are sometimes not co-localized with the biosynthetic genes, as was observed for streptolydigin, which inhibits the beta subunit of RNA polymerase (RNA PolB). The self-resistance *polB* gene is located outside of the cluster in the natural producer *Streptomyces lydicus*^21^.

### Expanding the repertoire of type I PKS gene clusters harboring one of the 14 validated self-resistance genes

To identify type I PKS clusters harboring one of the known self-resistance genes, we first performed a BLAST search of KS domains in all NCBI databases using 8 diverse KS domain sequences as queries (Figure 1). We ran antiSMASH 4.0 on all NCBI records harboring KS domains to detect clusters annotated as type I PKS and created a database of all proteins contained in these ∼30,000 clusters. We performed a BLAST search of all 14 experimentally validated self-resistance protein sequences against this protein database to identify which clusters harbor a putative self-resistance gene. Finally, we extracted all clusters in which the putative self-resistance gene is within 10 kB of a KS domain.

### Validation of the approach

18 clusters harbor an experimentally-validated self-resistance gene, including 9 type I PKS and type I PKS/NRPS hybrids. Our approach detected 8 out of 9, as the andrimid biosynthetic cluster was not annotated as a type I PKS by antiSMASH for unknown reasons, although it contains type I PKS KS domains. 7 of 8 clusters were identified using a distance cutoff of 10 kB between the KS and the resistance gene, whereas all 8 were identified using a 20 kB distance cutoff. Increasing the cutoff to 20 kB allowed the borrelidin biosynthetic cluster, which inhibits a threonyl-tRNA synthase, and whose gene cluster harbors a self-resistance copy of the threonyl-tRNA synthase gene 12 kB from a KS domain.

Of the ∼30,000 gene clusters annotated as type I PKSs by antiSMASH, 252 harbored a putative self-resistance gene within 10 kB of a KS domain (Table S2). A KS redundancy cutoff of 90% resulted in 190 non-redundant clusters. Clusters encoding the acetyl-CoA carboxylase beta subunit (ACCB) and isoleucyl-tRNA synthetases (Ile-tRNA synthetases) are the most abundant (Table 2). One explanation for that observation is that ACCB may be a biosynthetic enzyme providing the extender unit for the polyketide chain. Similarly, Ile-tRNA synthetase may be a biosynthetic enzyme involved in the biosynthesis of amino acids incorporated in the polypeptide moiety of a natural product. However, if the cluster harboring Ile-tRNA synthetase does not contain any NRPS modules, it is more likely that Ile-tRNA synthetase is a self-resistance protein. Similarly, for any putative self-resistance gene, if the corresponding protein is involved in the primary or secondary metabolism, it is difficult to assess whether its function within a cluster is to provide self-resistance or merely to encode a biosynthetic enzyme. Genes encoding non-metabolic enzymes, such as gyrase B, are less likely to be biosynthetic.

**Table 2.** Self-resistance genes vs. housekeeping genes: Number of PKS clusters harboring a putative self-resistance gene and more than one copy of that gene per nucleotide record.

| **N** | **Target Protein** | **Number of**<br>**Nonredundant**<br>**Clusters** | **Number of Clusters with**<br>**>1 Copy of Target per**<br>**Nucleotide Record** |
| --- | --- | --- | --- |
| 1 | 3-oxoacyl-[acyl-carrier-protein] synthase 1 (FabB/F) | 9 | 6 |
| 2 | Acetyl CoA carboxylase | 49 | 6 |
| 3 | Beta proteasome subunit | 8 | 3 |
| 4 | DNA polymerase sliding clamp | 4 | 1 |
| 5 | Elongation factor Tu | 17 | 5 |
| 6 | Enoyl-[acyl-carrier-protein] reductase [NADH] FabI | 1 | 0 |
| 7 | Gyrase B | 17 | 6 |
| 8 | Isoleucyl tRNA synthetase | 25 | 0 |
| 9 | Leucil-tRNA synthase | 6 | 0 |
| 10 | Methionine aminopeptidase | 1 | 0 |
| 11 | Ornithine carbamoyl transferase | 28 | 0 |
| 12 | Seryl-tRNA synthetase | 19 | 1 |
| 13 | Threonyl-tRNA synthetase | 6 | 1 |
| 14 | Tryptophanyl-tRNA synthase | 4 | 0 |

### Housekeeping copy of the putative self-resistance gene

We reasoned that self-resistance genes would often be a cluster-localized duplicate of a housekeeping gene located elsewhere in the genome, and that this genomic copy would not be resistant to the antibiotic. In the positive set of PKS clusters harboring validated self-resistance genes, only 4 out of 8 NCBI records containing BGCs were from complete or nearly-complete genomes. We manually examined those records for the co-occurrence of a housekeeping copy and we found one in all 4. The automated approach detected it in 3 records (Table 3): the sequence identity cutoff of 30% did not detect a second copy of the *ile-tRNA synthetase* in the genome of *Pseudomonas fluorescens*, which is only 23% identical to the self-resistance *ile-tRNA synthase* gene in the mupirocin biosynthetic cluster.

**Table 3.** Number of copies per genome in PKS clusters with experimentally validated self-resistance proteins.

| N | Natural Product | Target Protein | Genbank ID | Species | Number of copies in genome |
| --- | --- | --- | --- | --- | --- |
| 1 | thiolactomycin | 3-oxoacyl-[acyl-carrier-protein] synthase 1 (FabB/F) | KT282100 | *Salinispora pacifica* | 2 |
| 2 | thiotetramide | 3-oxoacyl-[acyl-carrier-protein] synthase 1 (FabB/F) | KT282101 | *Streptomyces afghaniensis* | 2 |
| 3 | salinosporamide | Beta proteasome subunit | CP000667 | *Salinispora tropica* | 2 |
| 4 | mupirocin | Isoleucyl tRNA synthetase | AF318063 | *Pseudomonas fluorescens* | 2 |

Only 29 of the 190 identified gene clusters are derived from complete genome sequences, and a second (putatively housekeeping) copy of the putative self-resistance gene was detected for 12 of those clusters (Table 2). The presence of a second housekeeping copy of the putative self-resistance gene is a useful feature by which to identify potential antibiotic producers, but sometimes only the self-resistance copy is sufficient for survival. For example, the rifamycin, streptolydiginm and streptovaricin producers *Amycolatopsis mediterranei, Streptomyces lydicus* and *Streptomyces spectabilis* harbor only one self-resistance copy of the RNA polymerase beta gene^21^. Furthermore, some genes exist in multiple copy number in bacterial genomes, and the copy number can differ from species to species^22^. As a result, a reference gene copy number per genome is difficult to establish. We applied the online tool ARTS to obtain a reference copy number, but ARTS output is limited to species in the Actinobacteria phylum. Thus, although informative, the presence of a housekeeping copy of the putative self-resistance gene is not a required characteristic of the antibiotic producer organisms.

### Phylogeny

We constructed a phylogenetic tree of all KS domains proximal to a putative self-resistance gene from the identified 190 PKS gene clusters harboring a putative self-resistance gene (Figure 2 and S1A). Two-thirds of the clusters originate from Actinobacteria, and the rest from Proteobacteria, Firmicutes and Cyanobacteria, and a small number of metagenomic sequences. We hypothesized that if a cluster was producing an antibiotic conferring selective advantage, that cluster would likely have a close homolog in another genome, also harboring a homologous self-resistance gene, either because of (a) horizontal gene transfer of the whole cluster, including the self-resistance gene, or (b) because the cluster will be conserved in closely related species. At least two examples support the latter: the borreledin BGC, which harbors a threonyl-tRNA synthetase self-resistance gene, is found in both *Streptomyces parvulus* (MIBiG ID BGC0000031) and in *Streptomyces rochei* (MIBiG ID BGC0001533). In addition, kalimantacin/batumin BGC, which harbors enoyl-[acyl-carrier-protein] reductase [NADH] FabI self-resistance gene is found in both *Pseudomonas fluorescens* (MIBiG ID BGC0001099) and *Pseudomonas batumici* (GenBank accession number JXDG01000003). Indeed, phylogenetic analysis revealed several clades in which the KS domains group with the putative self-resistance gene. Looking more closely at the architecture of the PKS clusters, we identified sets of homolog clusters: clusters which harbor the same number of modules and same number and set of domains. Interestingly, although there are 15 clusters harboring a gyrase B gene, the respective proximal KS domains have low sequence similarity. It is possible that phylogenetically distant type I PKS clusters target gyrase B, but the coverage of the sequence space is sparse and no homologs of these clusters have been discovered yet, or alternatively, there are no type I PKS clusters harboring a self-resistance gyrase B gene and the clusters identified are false positives.

**Figure 2.**
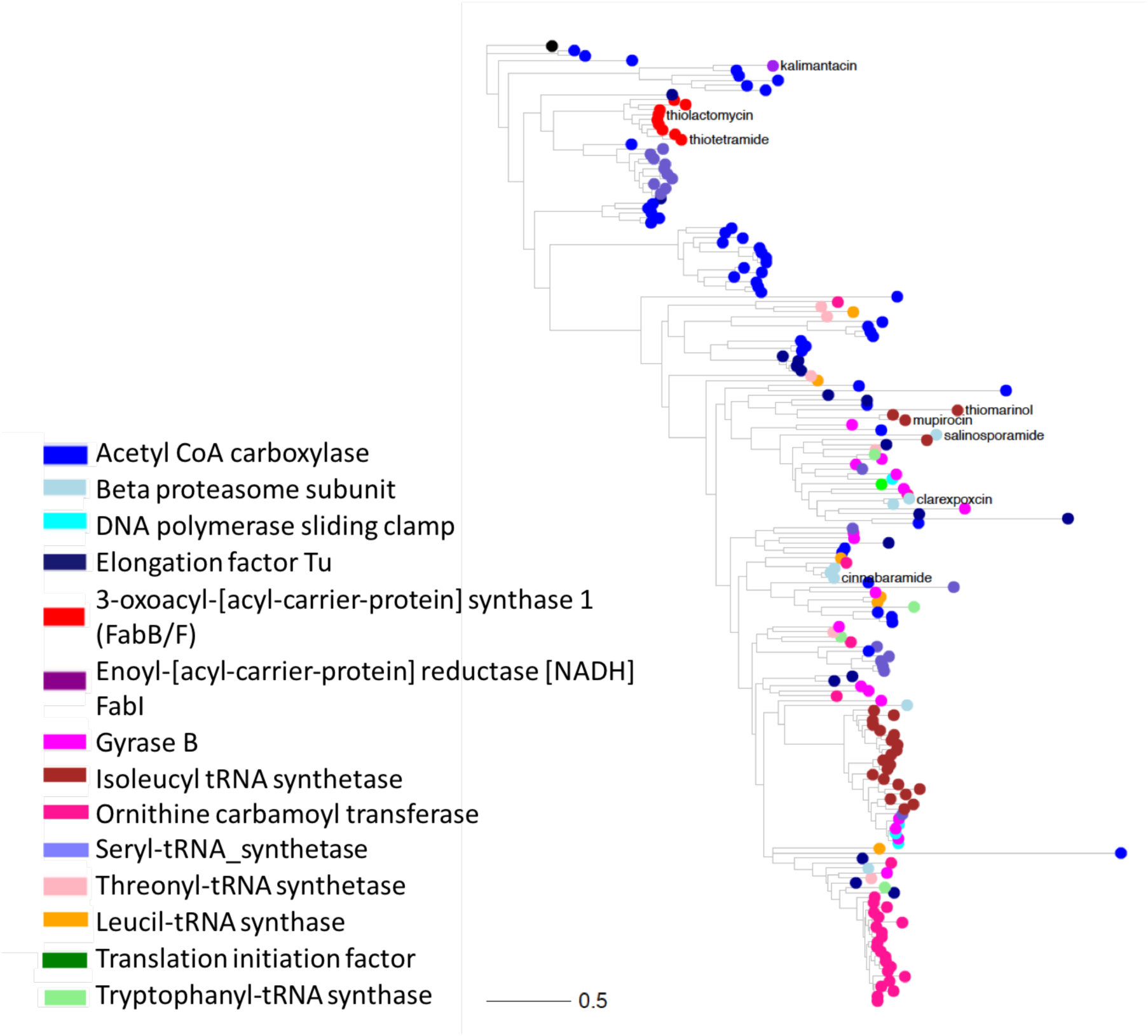
Phylogenetic tree of KS domains proximal to a self-resistance gene (190 KSs). Tree colored by presence of a putative self-resistance gene. PKS clusters harboring validated self-resistance genes (Table 1) are highlighted.

### Coevolution

We hypothesized that pairs of PKS gene clusters harboring self-resistance genes would exhibit a correlation between KS sequence identities and the corresponding putative self-resistance gene sequence identities. For example, if a gene cluster in one bacterial species gained a self-resistance gene, and the cluster subsequently evolved in other bacterial species, the KS pairwise similarities would track with the self-resistance gene pairwise identities. For each class of self-resistance gene, we calculated the Spearman correlation of percent identity of KS pairs and percent identity of putative self-resistance gene pairs (Table S1, Figure 3). Pairs of self-resistance proteins from certain classes (e.g., *fabB/F)* appear to have co-evolved with their proximal KS genes, suggesting possible co-evolution, whereas other classes (e.g., Gyrase B) do not show such co-evolutionary patterns (Figure 3). Most of the clusters harboring a *fabB/F* gene are similar to PKSs synthesizing known *fabB/F* inhibitors, thiolactomycin and thioteramide, suggesting a possible common origin.

**Figure 3.**
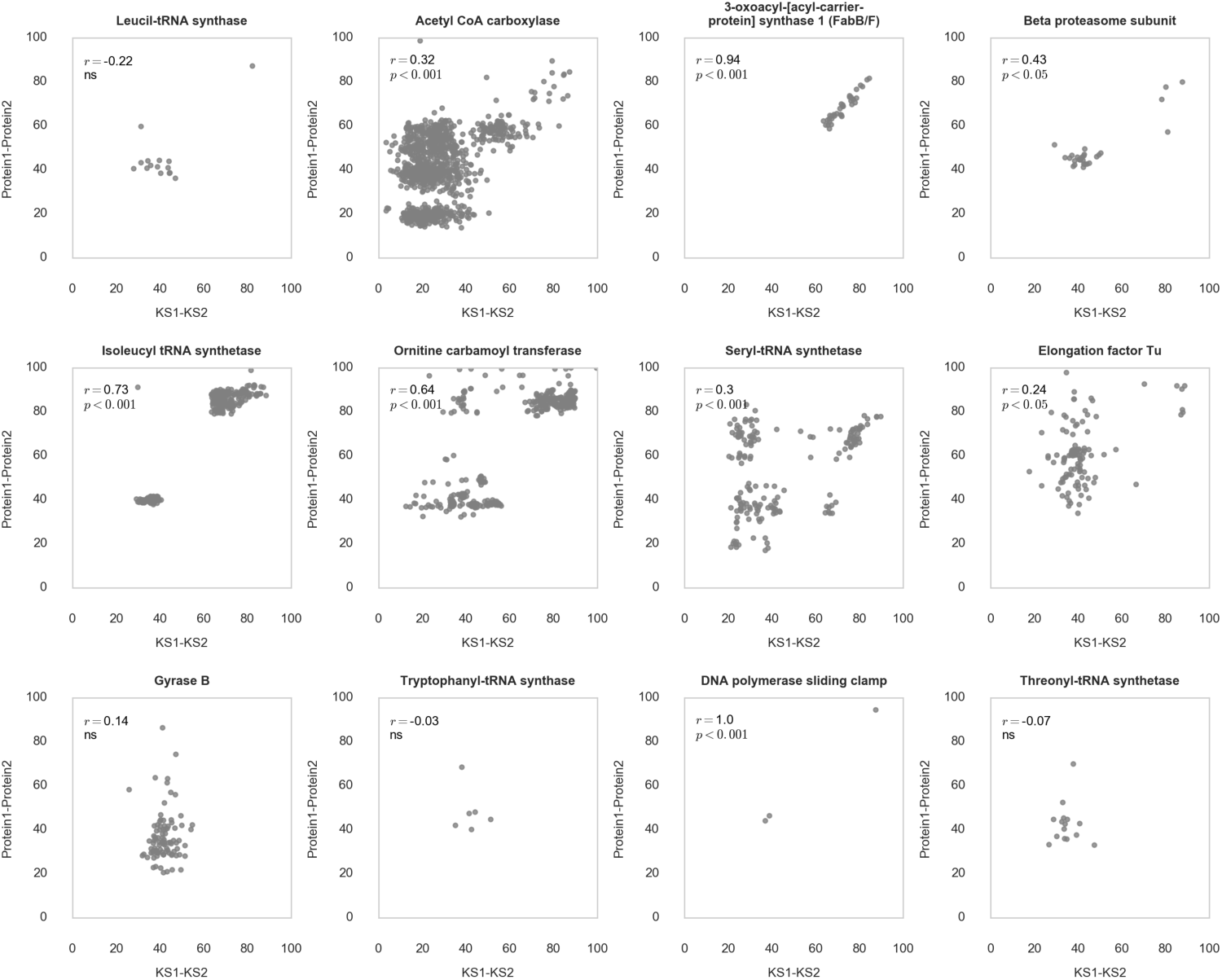
Coevolution of KS and putative self-resistance gene from the set of 14 experimentally validated targets. KS-self-resistance gene distance cutoff is 10 kB.

We measured correlation of KS pairwise identities and self-resistance gene pairwise identities for our newly-defined new self-resistance gene classes (Figure S2A and S2B). A correlation between KS identities and putative self-resistance gene identities suggests that the gene may have coevolved with the proximal KS gene. We refer to these correlations as co-evolution scores. Each class of putative self-resistance gene was assigned a co-evolution score (Figure S2A and S2B).

### Ranking clusters

To select the gene clusters most likely to encode a compound with antibiotic activity, we developed a ranking scheme based on (1) distance of the self-resistance gene to the biosynthetic KS gene, (2) presence of a “housekeeping” copy of the self-resistance gene, (3) coevolution score of the self-resistance gene to the biosynthetic KS gene, (4) presence of close homologs of the biosynthetic gene cluster in other genomes, and (5) ubiquity of the target. We attributed a score between 0 and 5 to each gene cluster; the scoring scheme is described in Table S1. As expected, most gene clusters from the positive set ranked highly (Table S2) with scores between 2 and 5, except for kalimantacin/batumin, which received a score of 0 because of distance of the self-resistance gene from the proximal KS, the lack of a housekeeping copy within the NCBI record, and the lack of close homologs in other microorganisms. We listed the top 20 ranking orphan clusters in Table 4.

**Table 4.** Top 20 ranked clusters harboring a putative self-resistance gene from the set of 14 experimentally validated targets.

| N | Target Name | GenBank ID | Cluster Number | Cluster Type | Species |
| --- | --- | --- | --- | --- | --- |
| 1 | 3-oxoacyl-[acyl-carrier-protein] synthase 1 (FabB/F) | CM002280 | cluster-3 | t1pks-terpene | *Streptomyces niveus* NCIMB 11891 |
| 2 | 3-oxoacyl-[acyl-carrier-protein] synthase 1 (FabB/F) | FOAZ01000002 | cluster-4 | butyrolactone-terpene-t1pks-otherks | *Streptacidiphilus jiangxiensis* |
| 3 | Beta proteasome subunit | BBUY01000002 | cluster-1 | bacteriocin-transatpks-t1pks-nrps | *Streptomyces* sp. NBRC 110611 |
| 4 | Beta proteasome subunit | QMEY01000002 | cluster-5 | phenazine-t1pks-nrps | *Sphaerisporangium* sp. LHW63015 |
| 5 | DNA polymerase sliding clamp | LZIX01000058 | cluster-1 | t1pks | *Mycobacterium* sp. E735 |
| 6 | Elongation factor Tu | QXCZ01000001 | cluster-1 | t1pks-otherks | *Streptomyces* sp. 3212.5 |
| 7 | Elongation factor Tu | NZ_LBDA02000011 | cluster-1 | t1pks-otherks | *Streptomyces malaysiense* |
| 8 | Isoleucyl tRNA synthetase | MBEU01000014 | cluster-1 | t1pks | *Mycobacterium* sp. IS-836 |
| 9 | Isoleucyl tRNA synthetase | LZLM01000005 | cluster-1 | t1pks-nrps | *Mycobacterium asiaticum* |
| 10 | Ornithine carbamoyl transferase | CP017593 | cluster-2 | t3pks-t1pks | *Mycobacterium tuberculosis* |
| 11 | Ornithine carbamoyl transferase | PEDJ01000075 | cluster-1 | t3pks-t1pks | *Mycobacterium marinum* |
| 12 | Ornithine carbamoyl transferase | LQOK01000037 | cluster-1 | t3pks-t1pks | *Mycobacterium bohemicum* |
| 13 | Ornithine carbamoyl transferase | LT841359 | cluster-1 | t3pks-t1pks | *Mycobacterium* sp. AFP003 |
| 14 | Ornithine carbamoyl transferase | LZKE01000216 | cluster-1 | t3pks-t1pks | *Mycobacterium* sp. E2327 |
| 15 | Ornithine carbamoyl transferase | NKRB01000001 | cluster-1 | t3pks-t1pks | *Mycobacterium kansasii* |
| 16 | Seryl-tRNA synthetase | AP010968 | cluster-2 | t1pks-otherks | *Kitasatospora setae* KM-6054 |
| 17 | Seryl-tRNA synthetase | CP015588 | cluster-3 | t1pks-butyrolactone | *Streptomyces alfalfae* |
| 18 | Seryl-tRNA synthetase | MTKC01000003 | cluster-2 | t1pks-otherks | *Streptomyces griseofuscus* |
| 19 | Seryl-tRNA synthetase | CP010849 | cluster-1 | t1pks-nrps | *Streptomyces cyaneogriseus* subsp. *noncyanogenus* |
| 20 | Seryl-tRNA synthetase | CM003601 | cluster-1 | t1pks | *Streptomyces anulatus* |

### ARTS comparison

In order to compare our approach to the ARTS platform, we ran ARTS on the top 20-ranked clusters. For approximately a third of these, the two approaches were in good agreement and ARTS also scored the clusters highly. However, for others of our top 20, such as cluster 3 of *Streptomyces niveus* NCIMB 11891 (NCBI accession CM002280), ARTS scored the cluster poorly. This is not surprising, since different metrics were used to calculate the final score. The two approaches are complementary and can be combined to evaluate clusters for antibacterial activity.

### Type I PKS catalog application

Our approach for searching type I PKS clusters harboring a putative self-resistance gene identified 190 non-redundant PKSs and PKS/NRPS hybrids. One way to visualize these gene clusters is a phylogenetic tree of their KS domains (Figure 2), including KS domains of the 9 known gene clusters that harbor validated self-resistance genes This phylogenetic map provides the opportunity to either study structural analogs of known clusters, or to explore orphan clusters for which the molecular structures of the encoded compounds are not known. Several clades lack any characterized cluster and could represent PKSs encoding new classes of polyketide antibiotics. For example, the Ile-tRNA inhibitor mupirocin belongs to a clade different that the one harboring most of the identified Ile-tRNA-synthetase-harboring clusters, which could encode a novel class of Ile-tRNA inhibitors. Similarly, a group of BGCs encode putative beta proteasome inhibitors, even though their proximal KS domains are not similar in sequence to that of salinosporamide synthase, a known proteasome inhibitor. These new BGCs could encode a new class of beta proteasomal inhibitors. We identified several homologs of thiolactomycin and thiotetramide BGCs (Figure 3, Table S2), which could be producing analogs of these compounds. Moreover, there are several clusters that harbor putative self-resistance genes encoding gyrase B, elongation factor Tu, and DNA polymerase sliding clamp, which were not previously reported as self-resistance genes in type I PKS clusters.

Ranking the discovered clusters should aid the prioritization of useful and biologically active natural products and should guide the refactoring of these biosynthetic gene clusters in tractable heterologous hosts. Clusters from non-actinobacterial origin, such as from Proteobacteria and Firmicutes, are attractive for expression in tractable non-actinobacterial heterologous hosts, such as *E. coli* or *B. subtilis*, respectively.

Most importantly, our approach provides hypotheses about the potential function and mechanism of action of the encoded compounds of all identified BGCs and could provide a useful resource for antibiotic discovery.

### Extending the approach to other antibacterial drug targets

Molecular targets for antibiotic discovery involved in DNA/RNA synthesis, protein synthesis, cell wall biosynthesis, and fatty acid biosynthesis have been characterized, but there is a need to characterize other known but underexplored targets^24^. We performed a literature search to identify known but underexplored antibacterial targets, hereafter referred to as “known” targets, by curating biosynthetic pathways of interest and the enzymes involved at each metabolic step, as well as antibacterial targets specific to multidrug resistant Gram-negative pathogens^25^. We searched MIBiG database for clusters harboring a potential self-resistance gene (Materials and Methods). We identified several BGCs with a known molecular target co-localized with the biosynthetic genes but not experimentally validated to be the self-resistance gene. Combining both approaches, we generated a list of 119 “known” targets from pathways of interest for antibacterial discovery. We searched our database of type I PKS gene clusters for these 119 target genes and identified 744 clusters using a 10 kB distance threshold between the KS and the putative self-resistance gene (Table S3, Figure S1B, Figure S2A). Out of 119 genes, 103 were observed to be co-localized with a type I PKS cluster. To prioritize which of these clusters might encode compounds with antibacterial activity, we ranked them using the above approach and identified 131 clusters with a score of 3 or higher. These high-ranking clusters harbor 34 putative self-resistance genes involved primarily in fatty acid biosynthesis, folate biosynthesis, cell wall biosynthesis, protein synthesis, protein degradation, DNA/RNA synthesis, and quinol/quinolone metabolism. Below we discuss a few interesting antibacterial targets and the clusters that harbor the potential self-resistance genes.

### Aminoacyl-tRNA synthetases as antibacterial targets

There are several known natural polyketide inhibitors of tRNA-synthetases: borrelidin (Thr-tRNA synthetase inhibitor), granaticin (Leucyl-tRNA synthase inhibitor), mupirocin A (Ile-tRNA synthetase inhibitor), reveromycin (Ile-tRNA synthetase inhibitor)^26^. Two of these, mupirocin and borrelidin, harbor a self-resistance ile- and thr-tRNA synthetase gene, respectively. However, problems with this class of antibacterials are (1) difficulty to penetrate the bacterial cell wall and enter the cell; (2) targets are structurally similar to eukaryotic tRNA-synthetase leading to human toxicity; (3) some of these compounds show cross-reactivity against targets other than tRNA synthetases. This makes tRNA-synthetase inhibitors a promising class of antibacterials, yet difficult to develop into clinically-approved therapies. Mupirocin is a naturally occurring Ile-tRNA synthetase inhibitor produced by *Pseudomonas fluorescens* and is the only aminoacyl-tRNA synthetase inhibitor used in clinic. We mined our type I PKS cluster database for all 20 aminoacyl-tRNA synthetases and identified several high-scoring clusters in Firmicutes, Proteobacteria and Actinobacteria, harboring a putative tRNA synthetase (Table S3).

A recently identified activity of bacterial Asp-tRNA and Glu-tRNA synthetase may provide yet-unexplored antibacterial targets. Some bacteria do not have Asn-tRNA and Gln-tRNA synthetase and instead use Asp-tRNA and Glu-tRNA synthetases to load the tRNA^Asn^ and tRNA^Gln^ with Asp and Glu respectively. The mischarged tRNA molecules are then converted to Asn-tRNA^Asn^ and Gln-tRNA^Gln^ and used in translation. These tRNA synthetases are structurally dissimilar from the canonical Asp-tRNA and Glu-tRNA synthetase^27^. Since they are also very different from eukaryotic tRNA synthetases, they are attractive antibacterial targets. Seven high-scoring clusters harbor an Asp-tRNA synthetase gene.

### Lipid A biosynthesis as antibacterial target pathway

With the increase in multidrug-resistant Gram-negative bacteria, there is imminent need for discovery of antibiotics with novel mechanisms of action. UDP-3-O-acyl-N-acetylglucosamine deacetylase (LpxC) is an enzyme that catalyzes the first committed step in lipid A biosynthesis^28^. Lipid A is a conserved component of lipopolysaccharides, which is a major component of the outer membrane of Gram-negative bacteria. LpxC is conserved in most Gram-negative bacteria (except in *Acinetobacter baumannii*) and it does not show homology to any mammalian protein and thus is considered an attractive target. Although drug-discovery efforts for LpxC inhibitors have been successful, no inhibitor has been approved to treat human infections to date^28^. The coevolution plot shows a strong correlation between pairwise KS and LpxC protein sequence identities (Figure S2A).

KS domains from all clusters encoding LpxC form a single clade (Figure S3). All 15 non-redundant clusters are annotated as Type I PKS by antiSMASH and their scores range from 2 to 5 (Table S3). They indeed have similar architecture, consisting of one fully reducing PKS module, and one or two additional KR genes. The lowest pairwise KS domains amino acid sequence identity is between CP003350 and AP013062 and is 73%. All clusters originate from the Proteobacteria phylum, mostly from the *Burkholderia* genus, but also from *Caballeronia, Frateuria, and Ralstonia* genera. The presence of 15 similar clusters across species, all of which harbor a LpxC gene, suggests that this cluster might be of evolutionary importance, and the target of the encoded molecule might be LpxC. However, it is also possible that the *lpxC* gene is a biosynthetic gene involved in the biosynthesis of the bacterial cell wall. Indeed, in mycobacteria, type I PKSs are implicated in the biosynthesis of surface-exposed lipopolisaccharides, which have roles in the mycobacterial virulence^29^. A few of the clusters harbor additional genes that might be involved in capsule polysaccharide biosynthesis, which supports the hypothesis that *lpxC* is a biosynthetic gene rather than a self-resistance gene. Further experiments are needed distinguish between these hypotheses.

### Proteases as antibacterial targets

Bacterial proteases are one set of underexploited antibacterial targets^23^. Proteases are protein degrading enzymes that play key roles in bacterial physiology, biochemistry, and pathogenicity. They have been shown to be druggable targets in mammals^23^, but currently there are no approved antimicrobial agents targeting bacterial proteases. There are four families of intracellular proteases in bacteria (ClpXP, Lon, HslUV and FtsH), and one periplasmic/secreted proteolytic complex (DegP). Bacterial proteasomes are mainly constrained to Actinobacteria phylum^30^. Several of these complexes have been explored as antibacterial targets. ClpP has been the most characterized and is the only complex for which natural product bacterial inhibitors have been found. Although cyclomarin, ecumicin and lassomycin are all active inhibitors of *M. tuberculosis* ClpP protease with high potency, further optimization of their pharmacological properties is required. The peptidyl boronate MG262 has been the only compound identified as a Lon inhibitor^31^. However, it is 2000-fold more potent against the human 20s proteasome, thus exhibiting off-target effects. There are no known HsIUV, FtsH or DegP inhibitors.

In order to identify novel protease inhibitors, we applied our approach in mining our database of proteins encoded in type I PKS clusters for ClpP, Lon, HsIUV, FtsH, and DegP and identified 54 clusters encoding at least one of them, including 10 clusters with scores of 3 or higher (Table S5.3, Table S5.4). Although most of the clusters are present in Proteobacteria, surprisingly, two clusters harboring a putative self-resistance *ftsH* gene are from fish: *Larimichthys crocea* (large yellow croaker) and *Scophthalmus maximus* (turbot). Both clusters consist of two PKS modules. Putative type I PKS clusters have been identified in metazoans, but only a few were functionally characterized: an iterative PKS encoding the yellow pigment of feathers in parakeets^32^, and a multi-modular PKS expressed during all larval stages in nematodes, possibly encoding a signaling molecule^33^. The newly identified clusters in fish could be false positives, because some antiSMASH-predicted domains span introns or intron/exon interfaces. However, it is possible that they might produce antibacterial compounds against bacterial proteases. These PKS clusters, together with the rest of the highly ranked clusters, provide a resource of potential inhibitors of bacterial proteases.

### Extending the pipeline to novel antibacterial drug targets

In order to identify novel antibacterial targets and the type I PKS clusters that may encode self-resistance genes, we hypothesized that any gene family that is known to be essential in bacteria could be an antibacterial target if it is in the proximity of a type I PKS cluster and if that clusters ranked highly in our scoring scheme. We mined our cluster database for clusters harboring a putative self-resistance gene from a set of 609 curated gene families that are essential in bacteria^34^. We added novel targets, for which there is no known inhibitor, from a manual literature search. This resulted in a list of 616 putative self-resistance genes. We identified 2470 non-redundant clusters that harbor one or more such genes within 10 kB of a core KS domain (Table S4, Figure S1C, Figure S2B). We selected all clusters that scored 3 or higher and compiled a table of all putative self-resistance genes in these clusters (Table S5, Figure 4), resulting in 89 unique putative self-resistance genes. Some of the identified self-resistance protein families, such as FabB/F and EF-Tu, already belong to the list of validated and “known” target families. The remaining 78 identified genes may be novel antibacterial targets.

**Figure 4.**
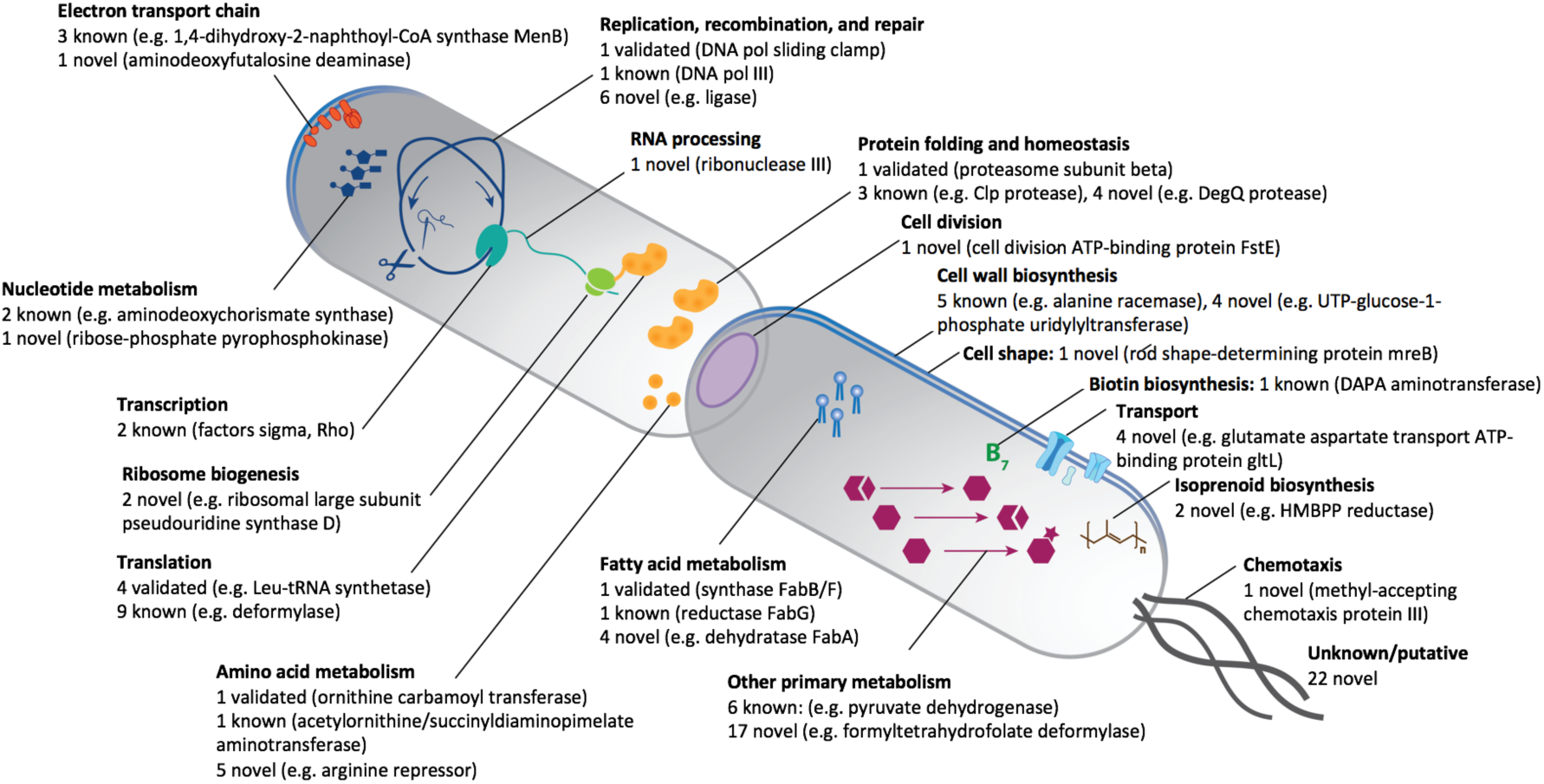
Antibacterial targets – validated, known, and putative. Clusters identified by mining 14 experimentally validated, 119 known, and 616 putative novel targets were ranked as described above. All clusters that scored higher than 3 were selected and a list of putative targets from these clusters was curated and categorized into several cellular processes. Examples of targets belonging to each category is shown below. The full list of validated, known, and novel targets is in Table S5.

We grouped the 89 target genes by pathway (Figure 4). Some of them belong to the well-established antibacterial target pathways: DNA replication, transcription and translation, nucleic acid and amino acid synthesis, fatty acid metabolism, cell wall biosynthesis and protein homeostasis.

Other genes belong to pathways that are essential for bacterial survival but are not directly targeted by any currently used antibiotic, such as cell division, cell shape maintenance, electron transport chain, RNA processing, ribosome biogenesis, chemotaxis, molecule transport, isoprenoid biosynthesis and other metabolic pathways. There is urgent need for new classes of antibiotics, and molecules targeting new pathways are a promising avenue for combating resistance against existing drugs. Some of these pathways are already being explored for the development of new antibacterials^35,36^. If molecules biosynthesized by identified clusters indeed target these pathways, this list would be a resource for bioprospecting in search for novel classes of antimicrobials. Many of these putative targets are hypothetical enzymes involved in primary or secondary metabolism and it is difficult to distinguish whether they provide self-resistance or are involved in the biosynthesis of the compound. However, many of the identified proteins are involved in non-metabolic pathways and are less likely to be biosynthetic.

To further prioritize clusters that are more likely to produce interesting compounds, we included additional information (Table S4). First, we indicated whether the target protein has a homolog in humans. When developing novel antibiotic compounds, toxicity is always an important consideration. Molecules that target bacterial proteins can exhibit off-target activity against their homologs in the eukaryotic host, as in the case of antibiotics targeting the ribosome^37,38^. For this reason, molecules targeting bacteria-specific proteins are more attractive drug candidates. We highlighted such targets by annotating proteins that do not have any known human homologs.

Second, we indicated whether the producer organism is Gram-negative. There are more challenges for antibiotic discovery for multi-drug resistant Gram-negative pathogens, since they have a double cell wall, which prevents antibiotic molecules from reaching their intracellular targets. Despite extensive efforts, there have been no new classes of antibiotics approved against Gram-negative pathogens in over fifty years^39^. Since our approach is based on the presence of a putative self-resistance gene within the cluster, it suggests that molecules produced by these clusters would target the host organism. Thus, clusters from Gram-negative bacteria are more likely to produce antibiotic molecules that are active against targets from other Gram-negative pathogens. Approximately one-third of targets in our list are from Gram-negative bacteria.

An interesting target pathway is protein folding and homeostasis. The proteasome is a known antibiotic target, and a self-resistance copy of its beta subunit is present in salinosporamide biosynthetic cluster^40^ and potentially other BGCs (Table 1). Moreover, several other proteins involved in protein homeostasis such as ClpX, Lon, as well as chaperone HtpG have been evaluated as potential antibacterial targets^23^. Our approach identified four other proteins encoded within BGCs which could represent promising antibacterial targets: proteases DegQ and FtsH, chaperone GroEL and the peptidyl-prolyl cis-trans isomerase PpiB that also acts as a chaperone. Bacterial proteases play essential roles in bacterial physiology and are exceptionally interesting targets, especially if they can be targeted by small molecules^23^. Chaperones represent another class of interesting targets. Despite the fact that high structural similarities between chaperones from bacteria and eukaryotes present a risk of toxicity, the development of antibiotics targeting bacterial chaperones is a promising avenue^41^. Our approach identified 14 highly ranked clusters harboring putative self-resistance genes *degQ, ftsH, groEL* and *ppiB* (Table S4). Further analysis of these clusters and their products could lead to new antibacterial molecules targeting these underexplored pathways.

## Conclusions

Despite recent advances in natural product discovery^42,43^, experimental identification of novel antimicrobial compounds and the corresponding BGCs lag behind progress in identifying gene clusters in newly-sequenced bacterial genomes. We used a bioinformatic approach to identify BGCs that may harbor self-resistance genes, revealing the target of the small molecule. Our technique provides guidance on the possible targets and modes of action of compounds encoded by BGCs. Our method can be generalized to gene clusters from any class of natural products, the biosynthesis of which involves a core enzyme, such as dimethylallyltryptophan synthase in alkaloids and trichodiene synthase in terpenes. It can be used to identify potential clusters not only from bacterial origin, but also from eukaryotic hosts. The approach can be used to prioritize gene clusters harboring potentially novel targets and encoding compounds with new mechanisms of action. This catalog of PKS clusters provides a resource for the discovery of new antibiotics with potentially novel targets. Many of the targets are specific for Gram-negative bacteria and lack human homologs, which makes them attractive antibacterial targets for future characterization.

## Notes

http://gvandova.com/publish/Data/

https://github.com/GerganaVandova/TargetMining

https://www.dropbox.com/sh/cjbcyyimgidl3q2/AAB_WF2I_K5FBkSA1JLarfOra?dl=0

### Competing Interest Statement

The authors have declared no competing interest.

