## Supplementary Information for "Identification of polyketide biosynthetic gene clusters that harbor self-resistance target genes"

### Supplementary Tables

#### Table S1. Ranking scheme.

| N | Parameters | Values of Parameters | Score |
|---|---|---|---|
| 1 | Distance to core enzyme | < 5 kB | 1 |
| 2 | Presence of close homologs | >1 homolog | 1 |
| 3 | Self-resistance gene copy number | >1 copy | 1 |
| 4 | Correlation coefficient | Spearman > 0.5, p-value < 0.05 | 1 |
| 5 | Target ubiquity | < 20 occurrences | 1 |

**Table S2. Catalog of PKS clusters identified through mining for 14 experimentally validated targets.** Table Clusters.14.10kb.xlsx is available online: http://gvandova.com/publish/Data/.

**Table S3. Catalog of PKS clusters identified through mining for 119 known targets.** Table Clusters.119.10kb.xlsx is available online: http://gvandova.com/publish/Data/.

**Table S4. Catalog of PKS clusters identified through mining for 616 novel targets.** Table Clusters.616.10kb.xlsx is available online: http://gvandova.com/publish/Data/.

**Table S5. List of experimentally validated, known, and novel targets found in proximity of the highly ranked clusters.**

Clusters identified by mining 14 experimentally validated, 119 known, and 616 putative novel targets were ranked as described above. All clusters that scored higher than 3 were selected and a list of putative targets from these clusters was curated. Targets were categorized into twenty distinct cellular processes. Targets with (+) are present in Gram-negative bacteria; for targets with (*) there are no human orthologs.

| N | Pathway | Validated Targets | Known Targets | Novel Targets |
|---|---|---|---|---|
| 1 | **Amino acid metabolism** | Ornithine carbamoyl transferase | Acetylornithine/succinyldiaminopimelate aminotransferase (P18335) | Arginine repressor (DEG10180483)<br>Dihydrodipicolinate synthase (DEG10180377)+<br>L-asparaginase II precursor (DEG10180443)+<br>Pyrroline-5-carboxylate reductase (DEG10180072)+<br>S-adenosylmethionine synthetase (DEG10180439)+ |
| 2 | **Biotin biosynthesis** | | Adenosylmethionine-8-amino-7-oxononanoate aminotransferase (P12995) | |
| 3 | **Cell division** | | | Cell division ATP-binding protein ftsE (DEG10180522)*+ |
| 4 | **Cell shape** | | | Rod shape-determining protein mreB (DEG10180486)*+ |
| 5 | **Cell wall biosynthesis** | | Alanine racemase (P0A6B4)<br>D-alanine-D-alanine ligase B (P07862)<br>UDP-3-O-acyl-N-acetylglucosamine deacetylase LpxC (P0A725)<br>UDP-N-acetylglucosamine 1-carboxyvinyltransferase MurA (P0A749)<br>UDP-N-acetylglucosamine-N-acetylmuramyl-(pentapeptide) pyrophosphoryl-undecaprenol N-acetylglucosamine transferase MurG (P17443) | 3-deoxy-manno-octulosonate cytidylyltransferase (DEG10180150)*+<br>Lipoprotein releasing system ATP-binding protein lolD (DEG10180185)<br>Probable transport ATP-binding protein msbA (DEG10180148)+<br>UTP-glucose-1-phosphate uridylyltransferase (DEG10180210)+ |
| 6 | **Chemotaxis** | | | Methyl-accepting chemotaxis protein III (DEG10180242)*+ |
| 7 | **Electron transport chain** | | 1,4-dihydroxy-2-naphthoyl-CoA synthase MenB (P0ABU0)<br>2-succinylbenzoate--CoA ligase MenE (P37353)<br>Na(+)-translocating NADH-quinone reductase subunit F (Q56584) | Aminodeoxyfutalosine deaminase (O86737_MQNX_STRCO) |
| 8 | **Fatty acid metabolism** | 3-oxoacyl-[acyl-carrier-protein] synthase 1 (FabB/F) | 3-oxoacyl-[acyl-carrier-protein] reductase FabG (P0AEK2) | (3R)-hydroxymyristoyl-[acyl carrier protein] dehydratase (DEG10180047)+<br>3-hydroxydecanoyl-[acyl-carrier-protein] dehydratase (DEG10180156)+<br>Acetyl-coenzyme A carboxylase carboxyl transferase subunit alpha (DEG10180052)+<br>Acyl carrier protein phosphodiesterase (DEG10180241) |
| 9 | **Isporenoid biosynthesis** | | | IspH protein (lytB) (DEG10180008)<br>Octaprenyl-diphosphate synthase (DEG10180474) |
| 10 | **Nucleotide metabolism** | | Aminodeoxychorismate synthase component 1 PabB (P05041) | Ribose-phosphate pyrophosphokinase (DEG10180204) |

| | | | | |
|---|---|---|---|---|
| | | | Aminodeoxychorismate synthase component 2 PanA (P00903) | |
| 11 | **Protein folding and homeostasis** | Proteasome subunit beta | ATP-dependent Clp protease ATP-binding subunit ClpX (P0A6H1)<br>Lon protease (P0A9M0)<br>Chaperone protein HtpG (P0A6Z3) | 60 kDa chaperonin GroEL (DEG10180584)[+]<br>ATP-dependent zinc metalloprotease FtsH (P0AAI3_ECOLI)[+]<br>Peptidyl-prolyl cis-trans isomerase B PpiB (DEG10180098)<br>Protease degQ precursor (DEG10180482)[+] |
| 12 | **Replication, recombination, repair** | DNA polymerase sliding clamp | DNA polymerase III subunit alpha (P10443) | Chromosomal replication initiator protein dnaA (DEG10180542)[+]<br>DNA gyrase subunit A (DEG10180351)<br>DNA ligase (DEG10180367)<br>Topoisomerase IV subunit A (DEG10180449)<br>Topoisomerase IV subunit B (DEG10180452) |
| 13 | **Ribosome biogenesis** | | | Ribosomal large subunit pseudouridine synthase D (DEG10180402)[+]<br>30S ribosomal protein S20 (DEG10180004) |
| 14 | **RNA processing** | | | Ribonuclease III (DEG10180397)[*+] |
| 15 | **Transcription** | | Transcription termination factor Rho (P0AG30)<br>ECF RNA polymerase sigma-E factor (P0AGB6) | |
| 16 | **Translation** | Elongation factor Tu<br>Isoleucyl-tRNA synthetase<br>Leucil-tRNA synthase<br>Seryl-tRNA synthetase | Alanyl-tRNA synthetase (DEG10180418)<br>Arginyl-tRNA synthetase (DEG10180318)<br>Aspartyl-tRNA synthetase (DEG10180315)<br>Elongation factor G (P0A6M8)<br>Elongation factor P (P0A6N4)<br>Glutamyl-tRNA synthetase (DEG10180366)<br>Peptide deformylase (P0A6K3)<br>Prolyl-tRNA synthetase (DEG10180054)<br>Translation initiation factor rubR1_TIF | |
| 17 | **Transport** | | | Glutamate aspartate transport ATP-binding protein gltL (DEG10180114)<br>Glutamate aspartate transport system permease protein gltK (DEG10180115)[+]<br>L-arabinose transport ATP-binding protein araG (DEG10180323)[+]<br>Oligopeptide transport ATP-binding protein oppD (DEG10180214) |
| 18 | **Transposition** | | | Insertion element IS1 4 protein insB (DEG10180163)[+] |
| 19 | **Other metabolic pathways** | | Dihydrolipoyl dehydrogenase (P99084)<br>Glutamine--fructose-6-phosphate aminotransferase [isomerizing] (P17169)<br>Glutamine synthetase (P9WN39)<br>Pyruvate carboxylase (Q9CHQ7)<br>Pyruvate dehydrogenase E1 component subunit alpha (P60089) | 1-deoxy-D-xylulose 5-phosphate synthase (DEG10180080)[+]<br>3-octaprenyl-4-hydroxybenzoate carboxy-lyase (DEG10180361)<br>Cysteine desulfurase (DEG10180388)[+]<br>Delta-aminolevulinic acid dehydratase (DEG10180069)<br>Dihydrolipoamide dehydrogenase (DEG10180029)<br>Formyltetrahydrofolate deformylase (DEG10180208)[+]<br>Glutamate-1-semialdehyde 2,1-aminomutase (DEG10180034)[+]<br>GTP cyclohydrolase I (DEG10180344)<br>GTP cyclohydrolase II (DEG10180222)<br>Phenylacetic acid degradation protein paaC (DEG10180238)<br>Phenylacetic acid degradation protein paaY (DEG10180240)[+]<br>Phospho-2-dehydro-3-deoxyheptonate aldolase, Trp-sensitive (DEG10180285) |

| | | | | |
|---|---|---|---|---|
| | | | | Serine hydroxymethyltransferase (DEG10180392)<br>SufA protein (DEG10180280)<br>Transketolase 1 (DEG10180438) |
| **20** | **Unknown/putative** | | | Hypothetical ABC transporter ATP-binding protein ydcT (DEG10180246)<br>Hypothetical ABC transporter ATP-binding protein ynjD (DEG10180300)<br>Hypothetical acetyltransferase yncA (DEG10180250)<br>Hypothetical oxidoreductase yciK (DEG10180220)<br>Hypothetical protein yaiN (DEG10180067)<br>Hypothetical protein ybiA (DEG10180132)<br>Hypothetical protein ydcN (DEG10180244)<br>Hypothetical protein ydgE (DEG10180267)<br>Hypothetical protein ydiI P77781 MENI ECOLI 1,4-dihydroxy-2-naphthoyl-CoA hydrolase (DEG10180281)<br>Hypothetical protein ydiO (DEG10180282)<br>Hypothetical protein ydjR (DEG10180296)<br>Hypothetical protein yfcH (DEG10180360)<br>Hypothetical protein ygaP (DEG10180415)<br>Hypothetical protein yliG (DEG10180135)<br>Hypothetical transport protein yeaN (DEG10180304)<br>Hypothetical transport protein yfaV (DEG10180356)<br>Protein ycgM (DEG10180198)<br>Protein ydjA (DEG10180302)<br>Protein yjgF (DEG10180597)<br>Probable ABC transporter ATP-binding protein yhbG (DEG10180475)<br>Putative beta-xylosidase (DEG10180061)<br>Putative methylase yhhF (DEG10180524) |

#### Supplementary Figures

**Figure S1. Phylogenetic trees of KS domains proximal to a self-resistance gene.** A) from the set of 14 validated targets; (B) from the set of 119 known targets, and (C) from the set 616 novel targets. High resolution image of all three phylogenetic trees can be found in Supplementary Materials.

A. B. C.

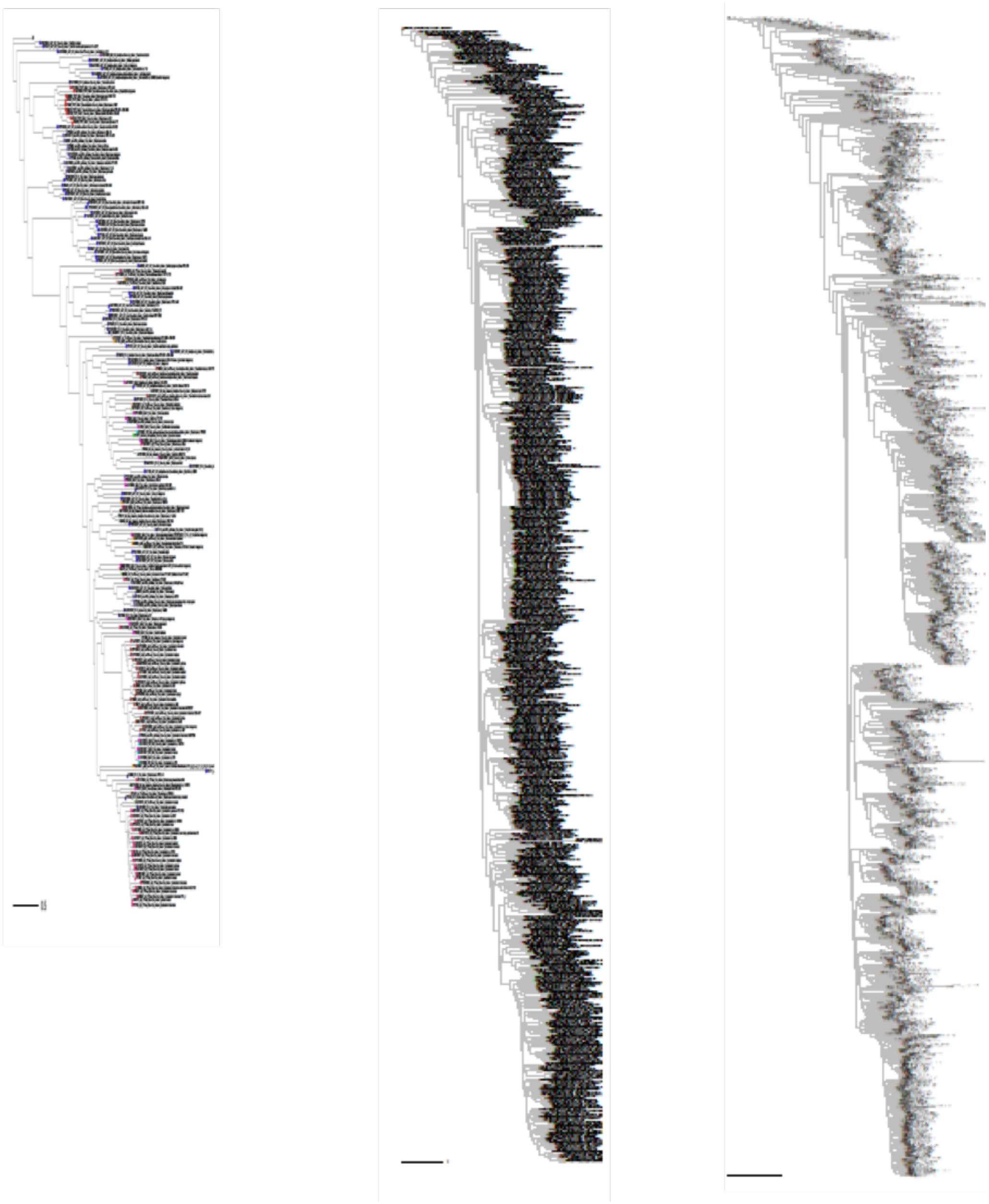

**Figure S2. Coevolution of KS and putative self-resistance gene.** KS-self-resistance gene distance cutoff is 10 kB. A) from the set of 119 known targets; B) from the set of 616 novel targets.

A.

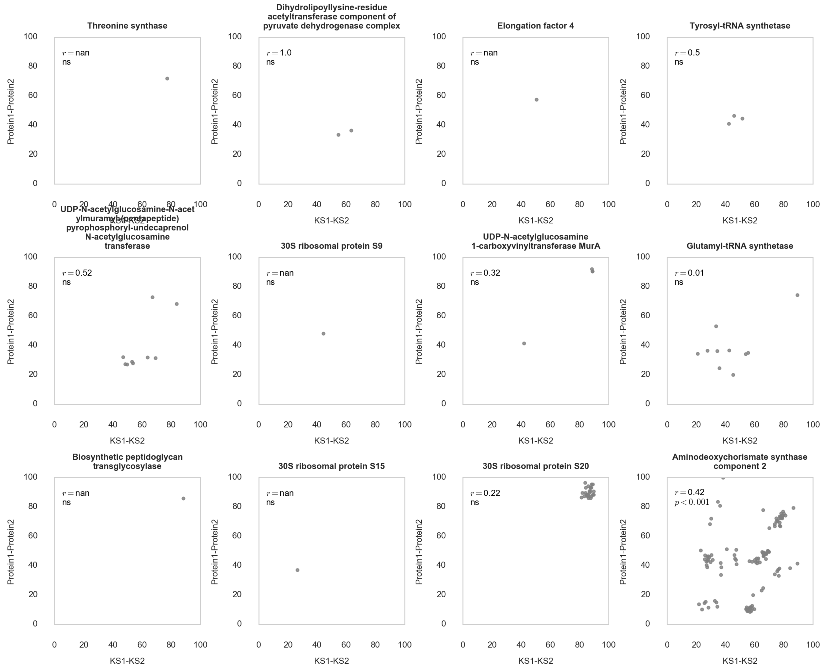

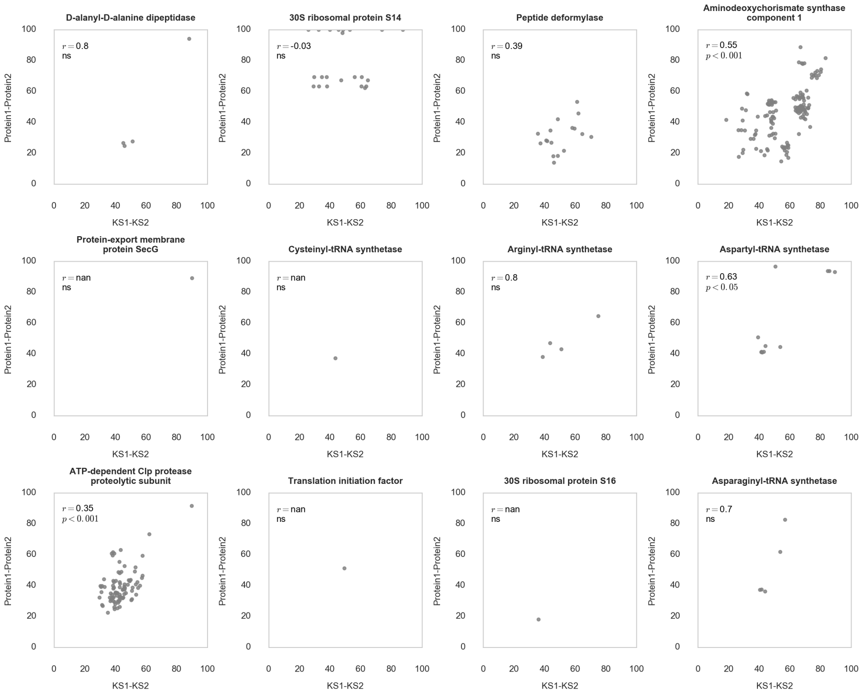
D-alanyl-D-alanine dipeptidase
$r = 0.8$
ns
30S ribosomal protein S14
$r = -0.03$
ns
Peptide deformylase
$r = 0.39$
ns
Aminodeoxychorismate synthase component 1
$r = 0.55$
$p < 0.001$
Protein-export membrane protein SecG
$r =$ nan
ns
Cysteinyl-tRNA synthetase
$r =$ nan
ns
Arginyl-tRNA synthetase
$r = 0.8$
ns
Aspartyl-tRNA synthetase
$r = 0.63$
$p < 0.05$
ATP-dependent Clp protease proteolytic subunit
$r = 0.35$
$p < 0.001$
Translation initiation factor
$r =$ nan
ns
30S ribosomal protein S16
$r =$ nan
ns
Asparaginyl-tRNA synthetase
$r = 0.7$
ns
Protein1-Protein2
KS1-KS2
0
20
40
60
80
100

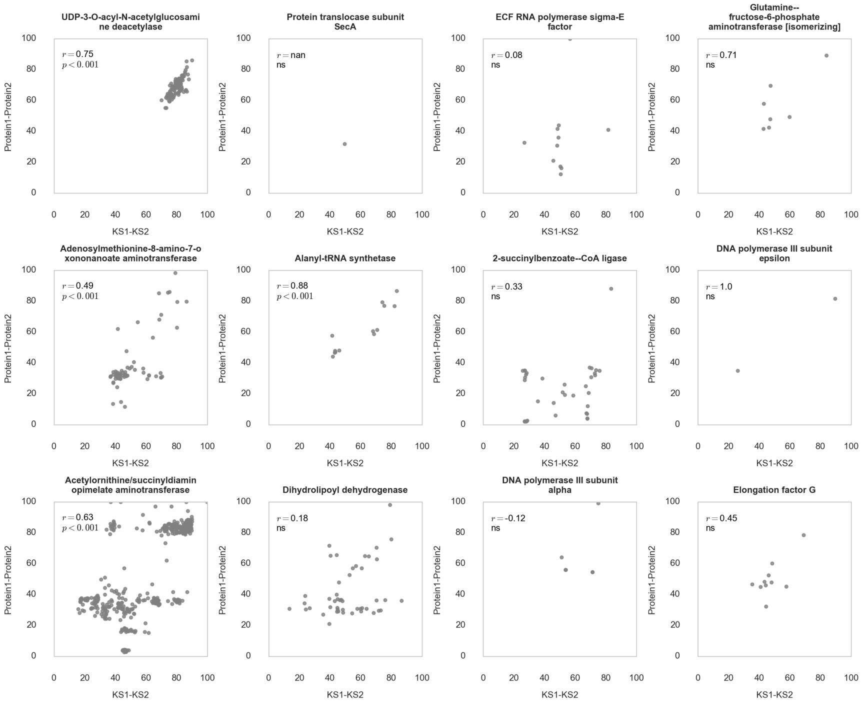
UDP-3-O-acyl-N-acetylglucosamine deacetylase
$r = 0.75$
$p < 0.001$
Protein translocase subunit SecA
$r =$ nan
ns
ECF RNA polymerase sigma-E factor
$r = 0.08$
ns
Glutamine--fructose-6-phosphate aminotransferase [isomerizing]
$r = 0.71$
ns
Adenosylmethionine-8-amino-7-oxononanoate aminotransferase
$r = 0.49$
$p < 0.001$
Alanyl-tRNA synthetase
$r = 0.88$
$p < 0.001$
2-succinylbenzoate--CoA ligase
$r = 0.33$
ns
DNA polymerase III subunit epsilon
$r = 1.0$
ns
Acetylornithine/succinyldiaminopimelate aminotransferase
$r = 0.63$
$p < 0.001$
Dihydrolipoyl dehydrogenase
$r = 0.18$
ns
DNA polymerase III subunit alpha
$r = -0.12$
ns
Elongation factor G
$r = 0.45$
ns
Protein1-Protein2
KS1-KS2
0
20
40
60
80
100

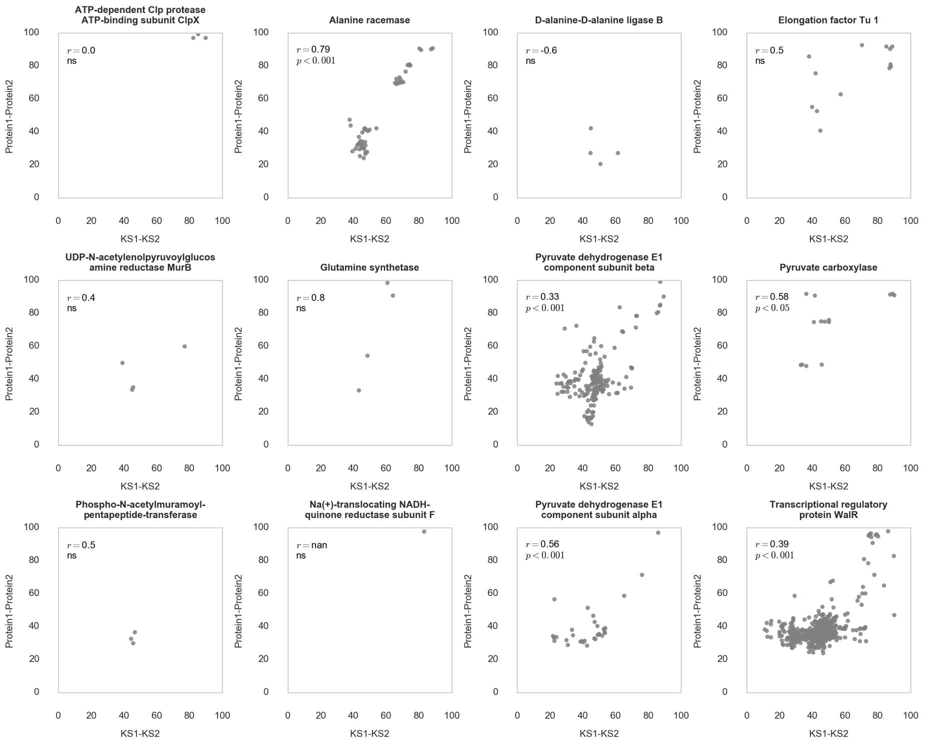
ATP-dependent Clp protease ATP-binding subunit ClpX
r = 0.0
ns
Alanine racemase
r = 0.79
p < 0.001
D-alanine-D-alanine ligase B
r = -0.6
ns
Elongation factor Tu 1
r = 0.5
ns
UDP-N-acetylenolpyruvoylglucos amine reductase MurB
r = 0.4
ns
Glutamine synthetase
r = 0.8
ns
Pyruvate dehydrogenase E1 component subunit beta
r = 0.33
p < 0.001
Pyruvate carboxylase
r = 0.58
p < 0.05
Phospho-N-acetylmuramoyl-pentapeptide-transferase
r = 0.5
ns
Na(+)-translocating NADH-quinone reductase subunit F
r = nan
ns
Pyruvate dehydrogenase E1 component subunit alpha
r = 0.56
p < 0.001
Transcriptional regulatory protein WalR
r = 0.39
p < 0.001
Protein1-Protein2
KS1-KS2

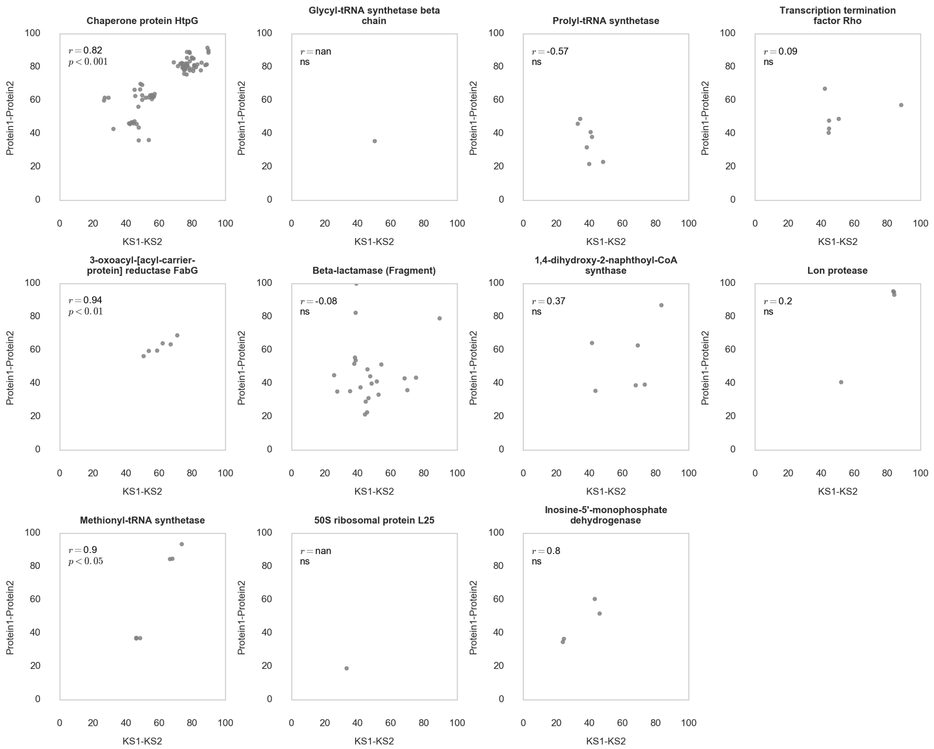
Chaperone protein HtpG
r = 0.82
p < 0.001
Glycyl-tRNA synthetase beta chain
r = nan
ns
Prolyl-tRNA synthetase
r = -0.57
ns
Transcription termination factor Rho
r = 0.09
ns
3-oxoacyl-[acyl-carrier-protein] reductase FabG
r = 0.94
p < 0.01
Beta-lactamase (Fragment)
r = -0.08
ns
1,4-dihydroxy-2-naphthoyl-CoA synthase
r = 0.37
ns
Lon protease
r = 0.2
ns
Methionyl-tRNA synthetase
r = 0.9
p < 0.05
50S ribosomal protein L25
r = nan
ns
Inosine-5'-monophosphate dehydrogenase
r = 0.8
ns
Protein1-Protein2
KS1-KS2

B.

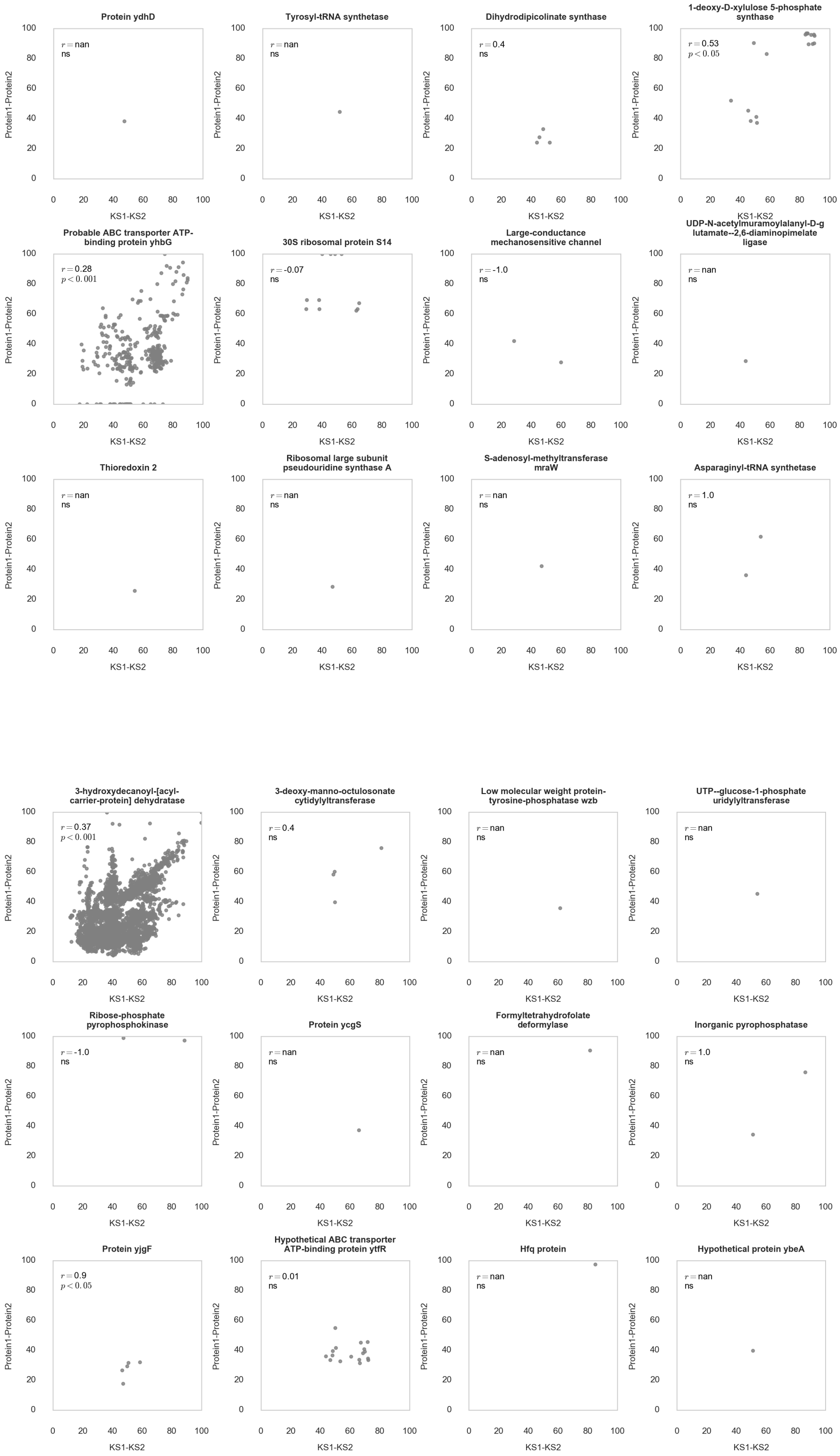
Protein ydhD
$r = $ nan
ns
Tyrosyl-tRNA synthetase
$r = $ nan
ns
Dihydrodipicolinate synthase
$r = $ 0.4
ns
1-deoxy-D-xylulose 5-phosphate synthase
$r = $ 0.53
$p < 0.05$
Probable ABC transporter ATP-binding protein yhbG
$r = $ 0.28
$p < 0.001$
30S ribosomal protein S14
$r = $ -0.07
ns
Large-conductance mechanosensitive channel
$r = $ -1.0
ns
UDP-N-acetylmuramoylalanyl-D-glutamate--2,6-diaminopimelate ligase
$r = $ nan
ns
Thioredoxin 2
$r = $ nan
ns
Ribosomal large subunit pseudouridine synthase A
$r = $ nan
ns
S-adenosyl-methyltransferase mraW
$r = $ nan
ns
Asparaginyl-tRNA synthetase
$r = $ 1.0
ns
3-hydroxydecanoyl-[acyl-carrier-protein] dehydratase
$r = $ 0.37
$p < 0.001$
3-deoxy-manno-octulosonate cytidylyltransferase
$r = $ 0.4
ns
Low molecular weight protein-tyrosine-phosphatase wzb
$r = $ nan
ns
UTP--glucose-1-phosphate uridylyltransferase
$r = $ nan
ns
Ribose-phosphate pyrophosphokinase
$r = $ -1.0
ns
Protein ycgS
$r = $ nan
ns
Formyltetrahydrofolate deformylase
$r = $ nan
ns
Inorganic pyrophosphatase
$r = $ 1.0
ns
Protein yjgF
$r = $ 0.9
$p < 0.05$
Hypothetical ABC transporter ATP-binding protein ytfR
$r = $ 0.01
ns
Hfq protein
$r = $ nan
ns
Hypothetical protein ybeA
$r = $ nan
ns

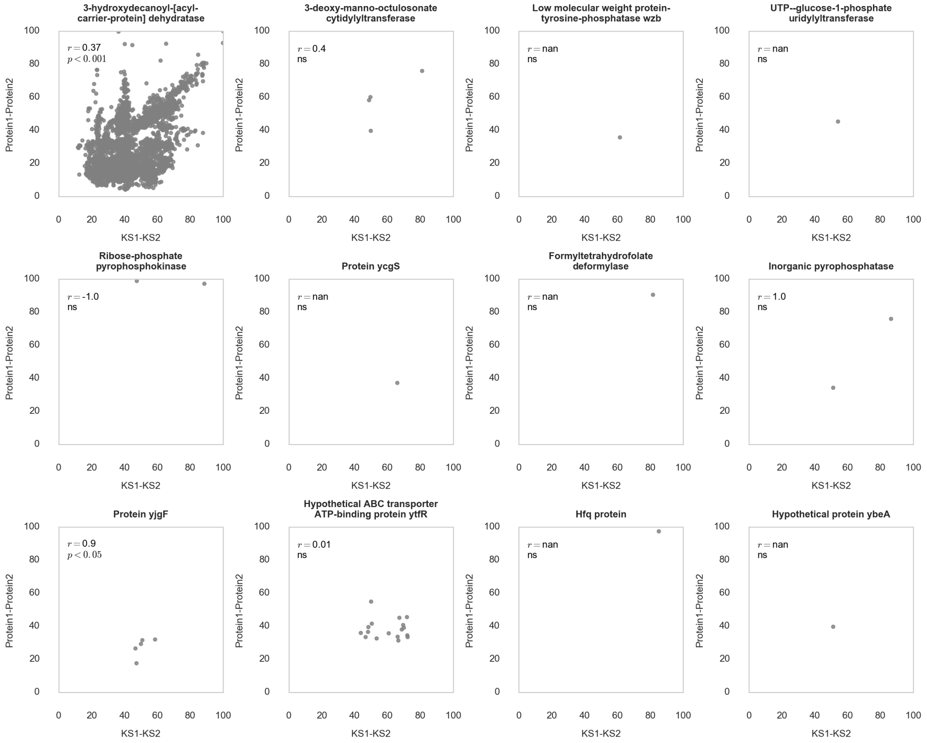
3-hydroxydecanoyl-[acyl-carrier-protein] dehydratase
$r = 0.37$
$p < 0.001$
3-deoxy-manno-octulosonate cytidylyltransferase
$r = 0.4$
ns
Low molecular weight protein-tyrosine-phosphatase wzb
$r = $ nan
ns
UTP--glucose-1-phosphate uridylyltransferase
$r = $ nan
ns
Ribose-phosphate pyrophosphokinase
$r = $ -1.0
ns
Protein ycgS
$r = $ nan
ns
Formyltetrahydrofolate deformylase
$r = $ nan
ns
Inorganic pyrophosphatase
$r = 1.0$
ns
Protein yjgF
$r = 0.9$
$p < 0.05$
Hypothetical ABC transporter ATP-binding protein ytfR
$r = 0.01$
ns
Hfq protein
$r = $ nan
ns
Hypothetical protein ybeA
$r = $ nan
ns
Protein1-Protein2
KS1-KS2

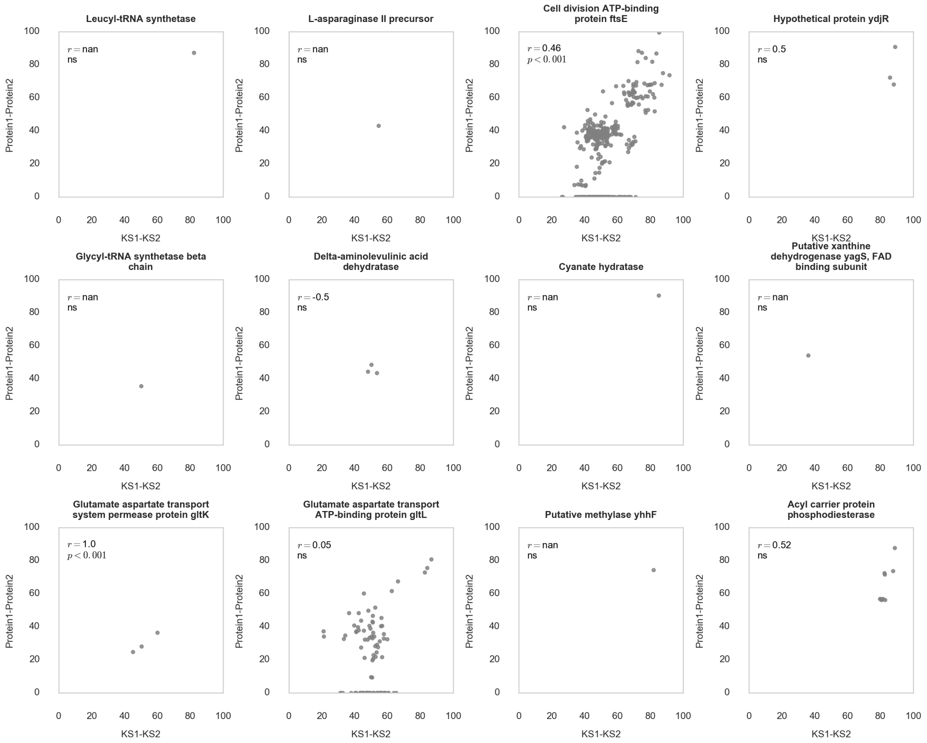
Leucyl-tRNA synthetase
$r = $ nan
ns
L-asparaginase II precursor
$r = $ nan
ns
Cell division ATP-binding protein ftsE
$r = 0.46$
$p < 0.001$
Hypothetical protein ydjR
$r = 0.5$
ns
Glycyl-tRNA synthetase beta chain
$r = $ nan
ns
Delta-aminolevulinic acid dehydratase
$r = $ -0.5
ns
Cyanate hydratase
$r = $ nan
ns
Putative xanthine dehydrogenase yagS, FAD binding subunit
$r = $ nan
ns
Glutamate aspartate transport system permease protein gltK
$r = 1.0$
$p < 0.001$
Glutamate aspartate transport ATP-binding protein gltL
$r = 0.05$
ns
Putative methylase yhhF
$r = $ nan
ns
Acyl carrier protein phosphodiesterase
$r = 0.52$
ns
Protein1-Protein2
KS1-KS2

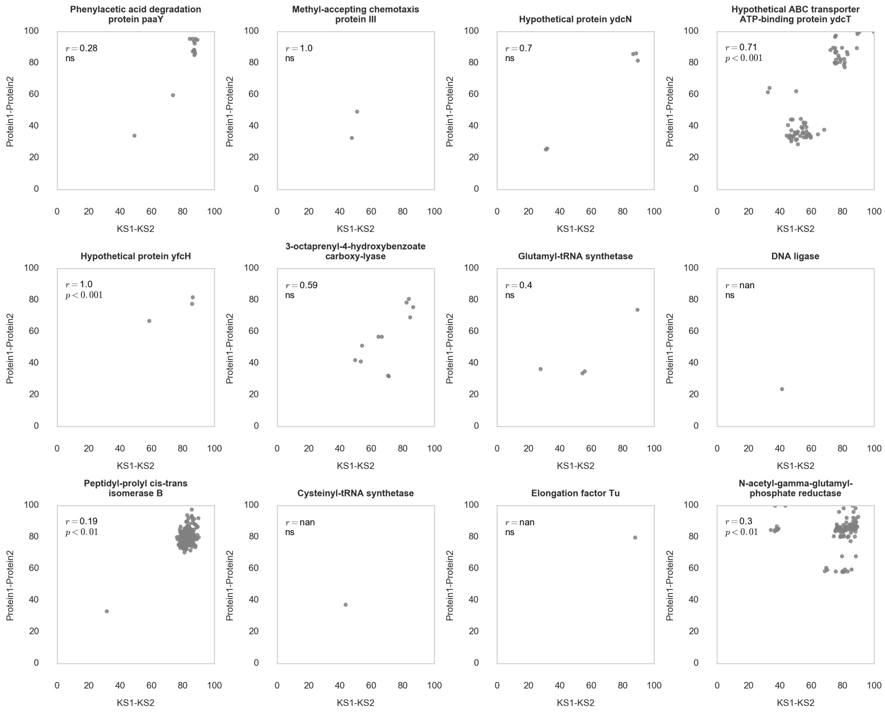
Phenylacetic acid degradation protein paaY
$r = 0.28$
ns
Methyl-accepting chemotaxis protein III
$r = 1.0$
ns
Hypothetical protein ydcN
$r = 0.7$
ns
Hypothetical ABC transporter ATP-binding protein ydcT
$r = 0.71$
$p < 0.001$
Hypothetical protein yfcH
$r = 1.0$
$p < 0.001$
3-octaprenyl-4-hydroxybenzoate carboxy-lyase
$r = 0.59$
ns
Glutamyl-tRNA synthetase
$r = 0.4$
ns
DNA ligase
$r =$ nan
ns
Peptidyl-prolyl cis-trans isomerase B
$r = 0.19$
$p < 0.01$
Cysteinyl-tRNA synthetase
$r =$ nan
ns
Elongation factor Tu
$r =$ nan
ns
N-acetyl-gamma-glutamyl-phosphate reductase
$r = 0.3$
$p < 0.01$
Protein1-Protein2
KS1-KS2

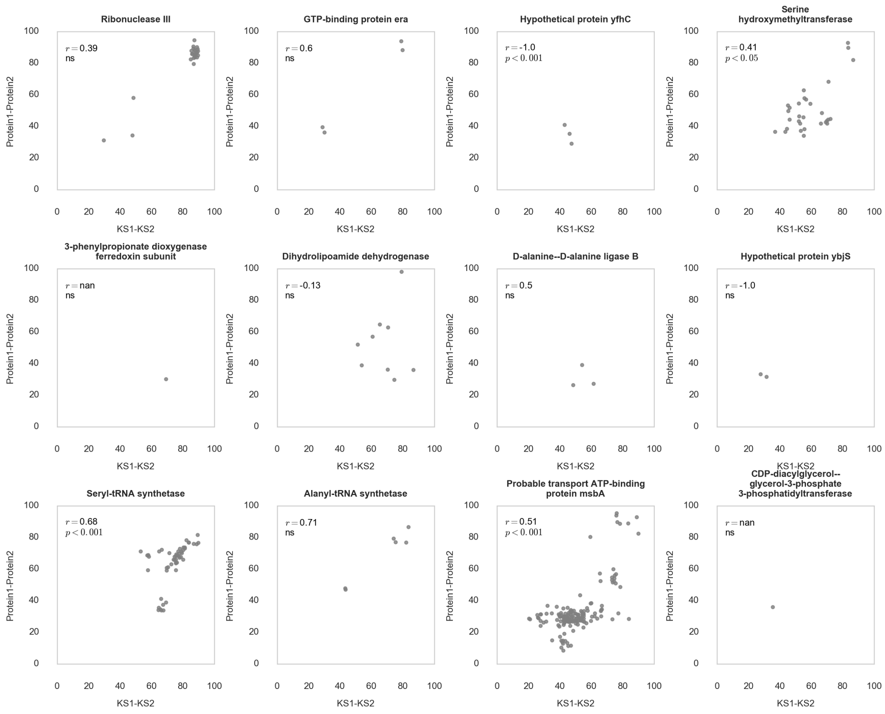
Ribonuclease III
$r = 0.39$
ns
GTP-binding protein era
$r = 0.6$
ns
Hypothetical protein yfhC
$r =$ -1.0
$p < 0.001$
Serine hydroxymethyltransferase
$r = 0.41$
$p < 0.05$
3-phenylpropionate dioxygenase ferredoxin subunit
$r =$ nan
ns
Dihydrolipoamide dehydrogenase
$r =$ -0.13
ns
D-alanine--D-alanine ligase B
$r = 0.5$
ns
Hypothetical protein ybjS
$r =$ -1.0
ns
Seryl-tRNA synthetase
$r = 0.68$
$p < 0.001$
Alanyl-tRNA synthetase
$r = 0.71$
ns
Probable transport ATP-binding protein msbA
$r = 0.51$
$p < 0.001$
CDP-diacylglycerol--glycerol-3-phosphate 3-phosphatidyltransferase
$r =$ nan
ns
Protein1-Protein2
KS1-KS2

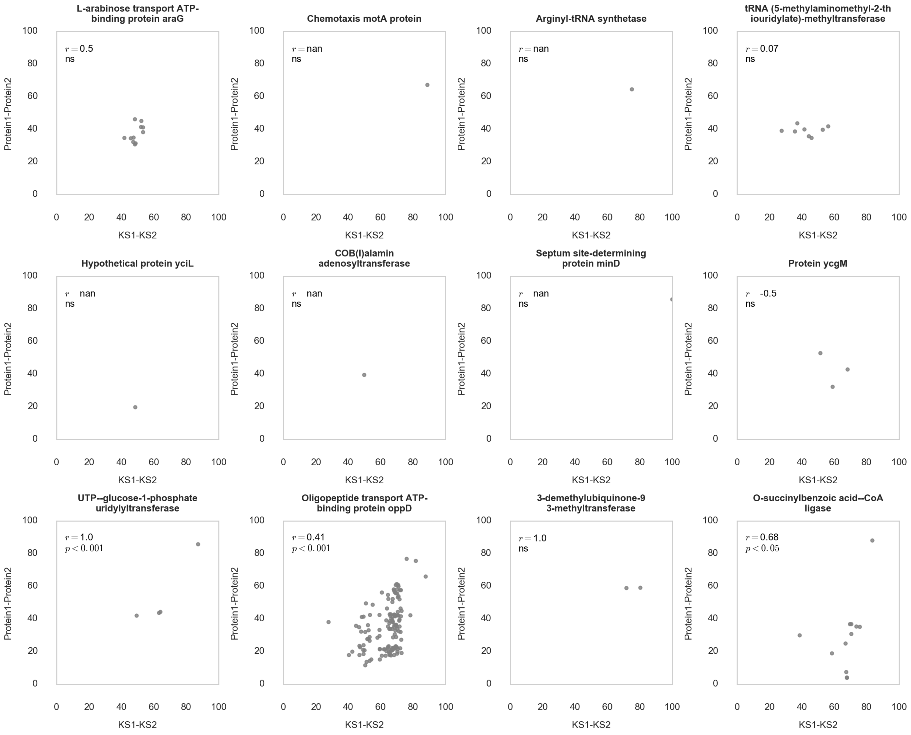
L-arabinose transport ATP-binding protein araG
r = 0.5
ns
Chemotaxis motA protein
r = nan
ns
Arginyl-tRNA synthetase
r = nan
ns
tRNA (5-methylaminomethyl-2-th iouridylate)-methyltransferase
r = 0.07
ns
Hypothetical protein yciL
r = nan
ns
COB(I)alamin adenosyltransferase
r = nan
ns
Septum site-determining protein minD
r = nan
ns
Protein ycgM
r = -0.5
ns
UTP--glucose-1-phosphate uridylyltransferase
r = 1.0
p < 0.001
Oligopeptide transport ATP-binding protein oppD
r = 0.41
p < 0.001
3-demethylubiquinone-9 3-methyltransferase
r = 1.0
ns
O-succinylbenzoic acid--CoA ligase
r = 0.68
p < 0.05
Protein1-Protein2
KS1-KS2

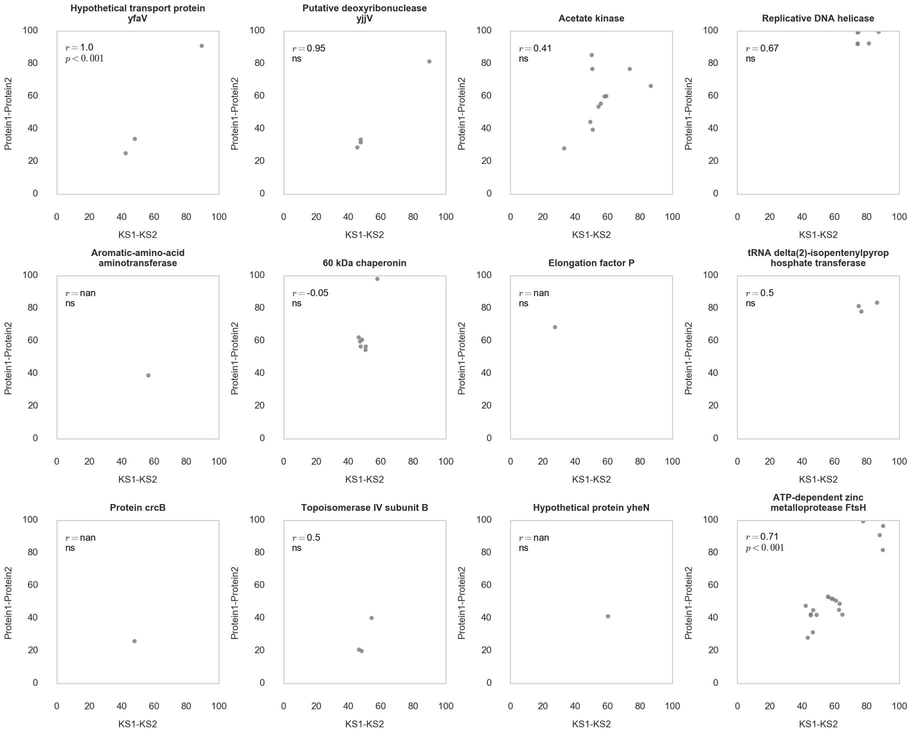
Hypothetical transport protein yfaV
r = 1.0
p < 0.001
Putative deoxyribonuclease yjjV
r = 0.95
ns
Acetate kinase
r = 0.41
ns
Replicative DNA helicase
r = 0.67
ns
Aromatic-amino-acid aminotransferase
r = nan
ns
60 kDa chaperonin
r = -0.05
ns
Elongation factor P
r = nan
ns
tRNA delta(2)-isopentenylpyrop hosphate transferase
r = 0.5
ns
Protein crcB
r = nan
ns
Topoisomerase IV subunit B
r = 0.5
ns
Hypothetical protein yheN
r = nan
ns
ATP-dependent zinc metalloprotease FtsH
r = 0.71
p < 0.001
Protein1-Protein2
KS1-KS2

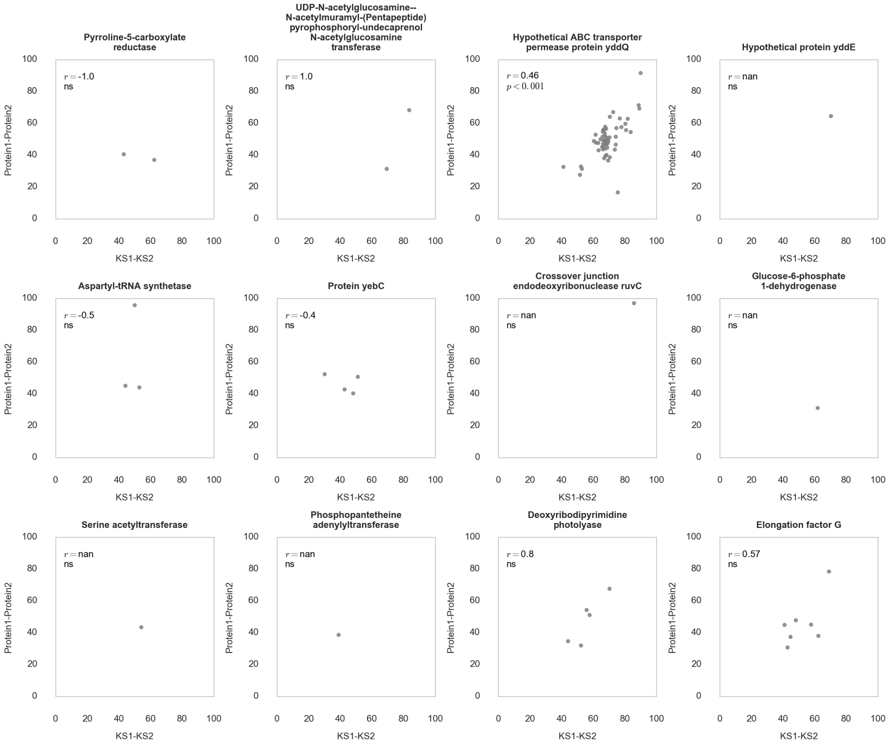
Pyrroline-5-carboxylate reductase
$r = -1.0$ ns
UDP-N-acetylglucosamine--N-acetylmuramyl-(Pentapeptide) pyrophosphoryl-undecaprenol N-acetylglucosamine transferase
$r = 1.0$ ns
Hypothetical ABC transporter permease protein yddQ
$r = 0.46$ $p < 0.001$
Hypothetical protein yddE
$r =$ nan ns
Aspartyl-tRNA synthetase
$r = -0.5$ ns
Protein yebC
$r = -0.4$ ns
Crossover junction endodeoxyribonuclease ruvC
$r =$ nan ns
Glucose-6-phosphate 1-dehydrogenase
$r =$ nan ns
Serine acetyltransferase
$r =$ nan ns
Phosphopantetheine adenylyltransferase
$r =$ nan ns
Deoxyribodipyrimidine photolyase
$r = 0.8$ ns
Elongation factor G
$r = 0.57$ ns
Protein1-Protein2
KS1-KS2

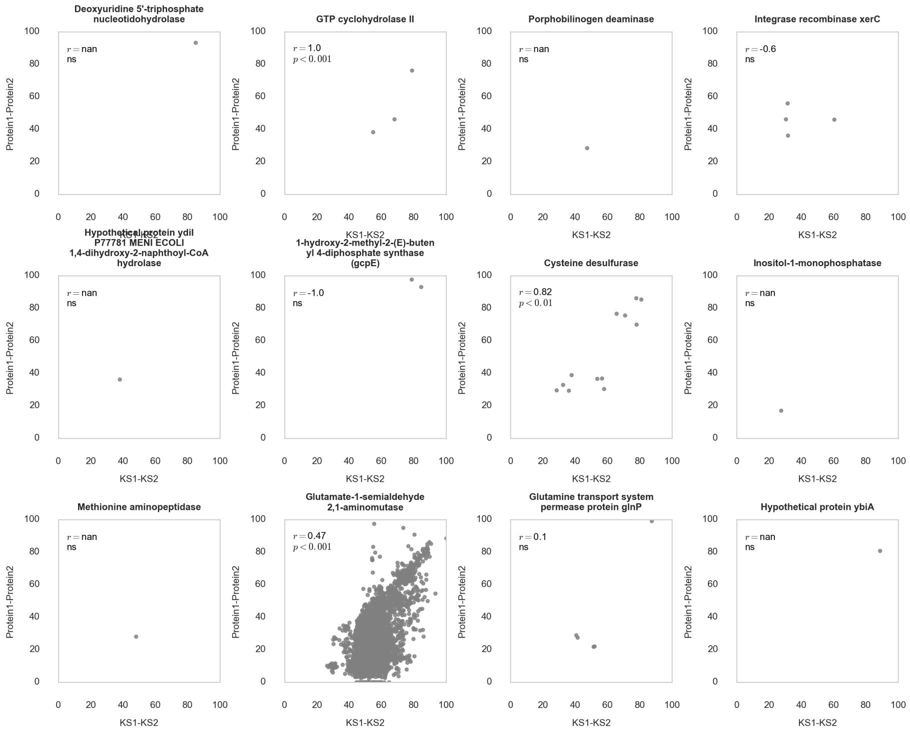
Deoxyuridine 5'-triphosphate nucleotidohydrolase
$r =$ nan ns
GTP cyclohydrolase II
$r = 1.0$ $p < 0.001$
Porphobilinogen deaminase
$r =$ nan ns
Integrase recombinase xerC
$r = -0.6$ ns
Hypothetical protein ydiI P77781 MENI ECOLI 1,4-dihydroxy-2-naphthoyl-CoA hydrolase
$r =$ nan ns
1-hydroxy-2-methyl-2-(E)-butenyl 4-diphosphate synthase (gcpE)
$r = -1.0$ ns
Cysteine desulfurase
$r = 0.82$ $p < 0.01$
Inositol-1-monophosphatase
$r =$ nan ns
Methionine aminopeptidase
$r =$ nan ns
Glutamate-1-semialdehyde 2,1-aminomutase
$r = 0.47$ $p < 0.001$
Glutamine transport system permease protein glnP
$r = 0.1$ ns
Hypothetical protein ybiA
$r =$ nan ns
Protein1-Protein2
KS1-KS2

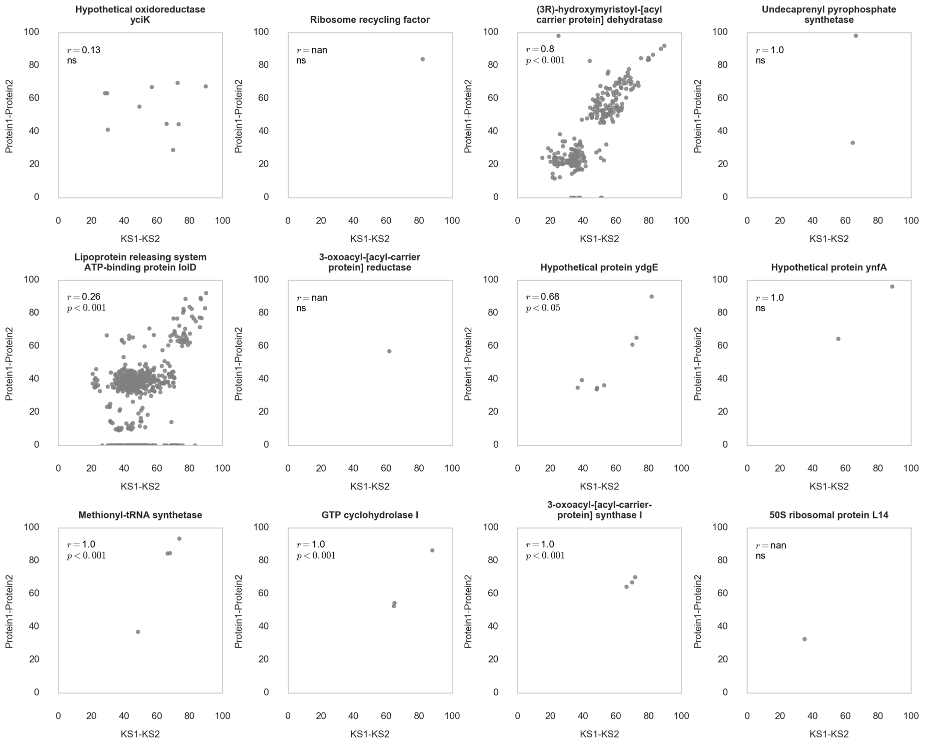
Hypothetical oxidoreductase yciK
r = 0.13
ns
Ribosome recycling factor
r = nan
ns
(3R)-hydroxymyristoyl-[acyl carrier protein] dehydratase
r = 0.8
p < 0.001
Undecaprenyl pyrophosphate synthetase
r = 1.0
ns
Lipoprotein releasing system ATP-binding protein lolD
r = 0.26
p < 0.001
3-oxoacyl-[acyl-carrier protein] reductase
r = nan
ns
Hypothetical protein ydgE
r = 0.68
p < 0.05
Hypothetical protein ynfA
r = 1.0
ns
Methionyl-tRNA synthetase
r = 1.0
p < 0.001
GTP cyclohydrolase I
r = 1.0
p < 0.001
3-oxoacyl-[acyl-carrier-protein] synthase I
r = 1.0
p < 0.001
50S ribosomal protein L14
r = nan
ns
Protein1-Protein2
KS1-KS2

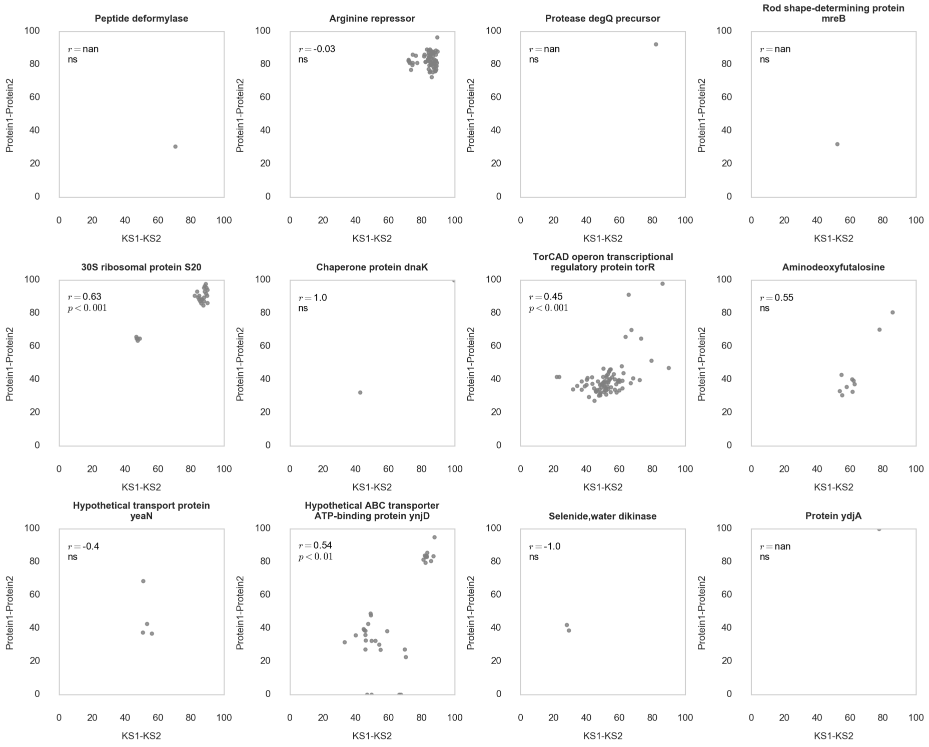
Peptide deformylase
r = nan
ns
Arginine repressor
r = -0.03
ns
Protease degQ precursor
r = nan
ns
Rod shape-determining protein mreB
r = nan
ns
30S ribosomal protein S20
r = 0.63
p < 0.001
Chaperone protein dnaK
r = 1.0
ns
TorCAD operon transcriptional regulatory protein torR
r = 0.45
p < 0.001
Aminodeoxyfutalosine
r = 0.55
ns
Hypothetical transport protein yeaN
r = -0.4
ns
Hypothetical ABC transporter ATP-binding protein ynjD
r = 0.54
p < 0.01
Selenide,water dikinase
r = -1.0
ns
Protein ydjA
r = nan
ns
Protein1-Protein2
KS1-KS2

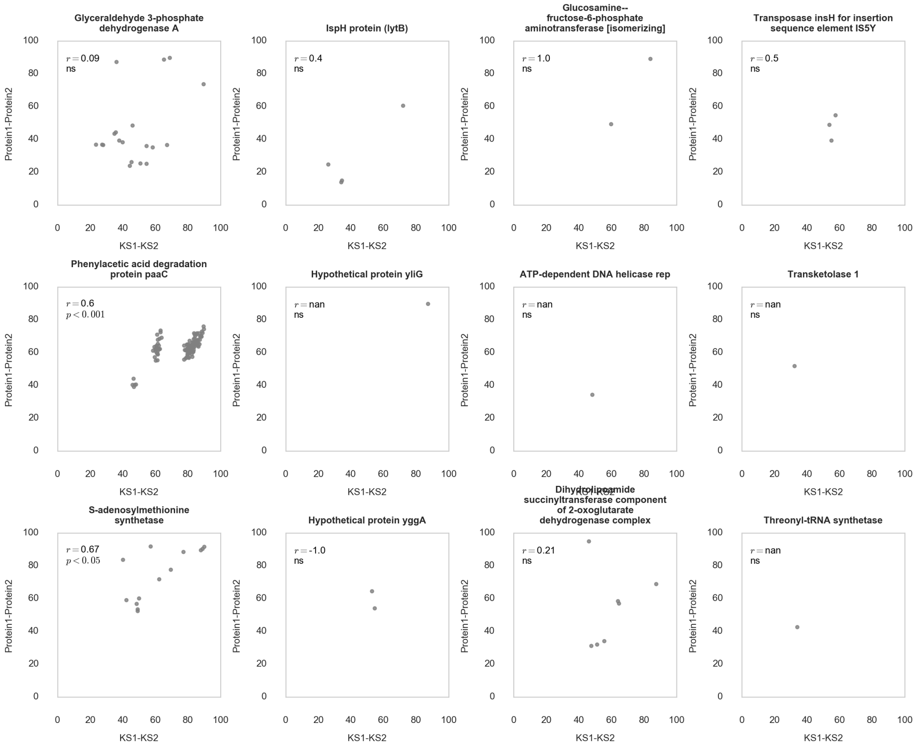
Glyceraldehyde 3-phosphate dehydrogenase A
$r = 0.09$
ns
IspH protein (lytB)
$r = 0.4$
ns
Glucosamine--fructose-6-phosphate aminotransferase [isomerizing]
$r = 1.0$
ns
Transposase insH for insertion sequence element IS5Y
$r = 0.5$
ns
Phenylacetic acid degradation protein paaC
$r = 0.6$
$p < 0.001$
Hypothetical protein yliG
$r =$ nan
ns
ATP-dependent DNA helicase rep
$r =$ nan
ns
Transketolase 1
$r =$ nan
ns
S-adenosylmethionine synthetase
$r = 0.67$
$p < 0.05$
Hypothetical protein yggA
$r = -1.0$
ns
Dihydrolipoamide succinyltransferase component of 2-oxoglutarate dehydrogenase complex
$r = 0.21$
ns
Threonyl-tRNA synthetase
$r =$ nan
ns
Protein1-Protein2
KS1-KS2

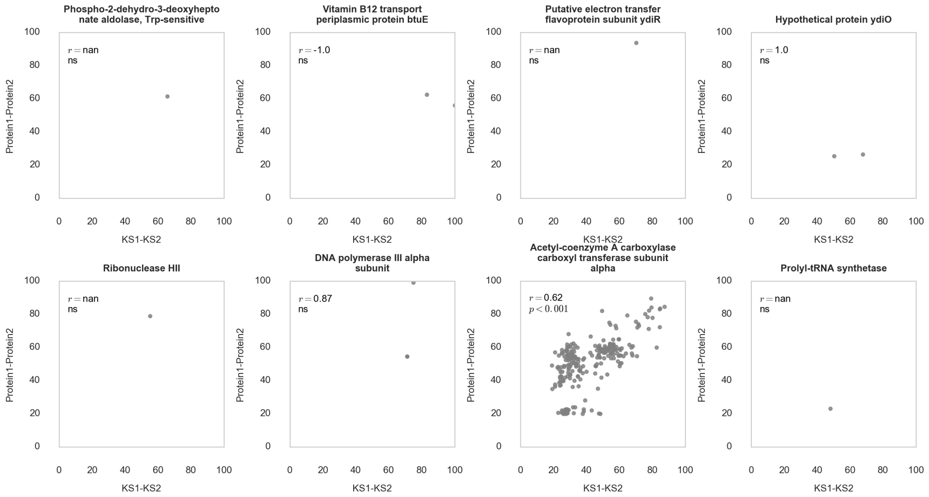
Phospho-2-dehydro-3-deoxyhepto nate aldolase, Trp-sensitive
$r =$ nan
ns
Vitamin B12 transport periplasmic protein btuE
$r = -1.0$
ns
Putative electron transfer flavoprotein subunit ydiR
$r =$ nan
ns
Hypothetical protein ydiO
$r = 1.0$
ns
Ribonuclease HII
$r =$ nan
ns
DNA polymerase III alpha subunit
$r = 0.87$
ns
Acetyl-coenzyme A carboxylase carboxyl transferase subunit alpha
$r = 0.62$
$p < 0.001$
Prolyl-tRNA synthetase
$r =$ nan
ns
Protein1-Protein2
KS1-KS2

#### Supplementary Materials

All supplementary data is published online: http://gvandova.com/publish/Data/. This resource contains:

1. FASTA files of validated, known, and novel targets, which were used to mine NCBI databases;
2. High resolution phylogenetic trees for each target dataset; they can also be found in https://www.dropbox.com/sh/cjbcyyimgidl3q2/AAB_WF2I_K5FBkSA1JLarfOra?dl=0.
3. Catalogues of PKS and PKS/NRPS clusters identified through mining for experimentally validated (Table S2. Clusters.14.10kb), known (Table S3. Clusters.119.10kb) and putative novel (Table S4 Clusters.616.10kb) targets;
4. Individual coevolution plots for each putative self-resistance gene.
